# Intense oocyte competition builds the female reproductive reserve in mice

**DOI:** 10.1101/2025.01.30.635812

**Authors:** Yan Zhang, Yingnan Bo, Kaixin Cheng, Lu Mu, Jing Liang, Lingyu Li, Xindi Hu, Ge Wang, Kaiying Geng, Xuebing Yang, Wenji Wang, Longzhong Jia, Xueqiang Xu, Jingmei Hu, Chao Wang, Yuwen Ke, Guoliang Xia, Hua Zhang

**Author notes:** Corresponding author: Hua Zhang.

## Abstract

During the process of ovariogenesis, more than two-thirds of germ cells are sacrificed to enhance the quality of the remaining oocytes. But the detail process of selection is not completely understood in mammals. Here, by creating a high resolution 4- dimensional ovariogenesis imaging system, we recorded the progress of oocyte fate determination in live mouse ovaries, and discovered an oocyte competition mechanism that determines the survival of oocytes. We found that dominant oocytes catch and absorb the cell debris from sacrificed oocytes to enrich their cytoplasm for survival. Single-cell sequencing demonstrated that this competition follows a classic autophagy-driven model. Blocking competition by suppressing autophagy resulted in an enlarged pool of survived oocytes. However, these oocytes lacking competition were unable to fully develop and contribute to fertility. Our study suggests that mammals have evolved a unique intensive competition system to enhance oocyte quality, necessary to support their long reproductive lifespan.

## Introduction

In sexually reproductive organisms, life begins with the combination of oocyte and sperm, and a successful fertilization requires high-quality gametes especially the oocyte^1^. Therefore, females developed a complex system for improving oocyte quality by sacrificing part of the oocytes to nurse the selected oocytes during ovariogenesis ^2,3^. These selections are believed to be a highly programmed and conserved process in both invertebrate and vertebrate species ^4^. To understand this process, a large body of studies has focused on oogenesis in invertebrate model species, and the classic model of oocyte selection is derived from *Drosophila*. In this model, germ cells rapidly divide and form germline cyst structures in which sister oocytes are connected with incompletely divided cytoplasm ^3,5^. Organelle-enriched cytoplasm is then orderly transported from nursing germ cells to the dominant one through cytoplasmic bridges, which forms the winner-oocyte for creating new life ^4,6,7^. In the model, oocyte connections and communications through intercellular bridges (IBs) in the cysts are believed to be indispensable for the determination of oocyte fate ^3,4^.

The life cycle of oocytes in mammals is much complex than that in invertebrate species. During ovariogenesis in the fetal and neonatal periods, a small portion of germ cells is selected through sacrifice of the other 2/3 of total germ cells to construct a non-renewable ovarian reserve ^8,9^. After the selection, oocytes must survive for more than 1 year in rodents and around 50 years in humans to maintain the normal reproductive lifespan ^10^. By tracing the oocyte development using a non-specific inducible cell labeling strategy in mice, Lei et al., showed that the organelle-enriched cytoplasm is transported from nursing germ-cells to dominant cells through IBs in cysts, which finally decides the survival oocytes to construct the ovarian reserve ^2,8,11^. Recently, Niu et al. described the detail model of cytoplasm exchanges in mouse ovarian cysts and dissected the inner molecular mechanisms in this process by single cell RNA sequencing. These studies provided important information to understand the model of early oocyte selection and development and demonstrated that forming oocyte cysts and the communications of sister oocytes in cysts are essential for their fate determination ^12^. All of these studies highlighted that the early selection of oocytes follows a conserved model in mammals as in flies.

However, the fertile phenotype in the *Tex14* knockout mouse model, in which females lacked IBs, suggested that cyst structure might not be indispensable for oocyte fate determination and the cyst-dependent germ cell model might not be the only mechanism for the oocyte selection in mice ^13,14^. Considering that the longevity of mammalian oocytes is much longer than that in invertebrate species, whether the mammals have evolved any other specific strategies of oocyte selection to boost the quality of survived oocytes for fitting their long lifespan of reproduction remains elusive. In the meantime, although there is a wealth of evidence from in vivo and in vitro studies indicating the indispensable role of cyst structures in selecting and determining the fate of oocytes ^4^, there has been a lack of long-term, detailed live observation and tracking of early oocyte development within the ovary. This has left us with limited knowledge of the specific mechanisms and related cell behaviors involved in oocyte selection.

Over the past decade, there has been a growing body of evidence indicating that cell competition, whereby higher fitness cells outcompete and eliminate lower fitness cells, plays a significant role in various biological events during mammalian embryonic development ^15,16^. This selection mechanism is characterized by a range of intense cellular behaviors, including the engulfment of loser cells by winner cells to enhance their own development ^17,18^. Interestingly, the key regulators of cell competition, such as autophagy signaling ^19^ and *Tp53* ^20–22^, have also been found to be involved in mammalian oocyte selection, providing indications that cell competition may also occur during mammalian ovariogenesis. Nevertheless, further experimental evidence is needed to definitively confirm this idea.

In our study, we utilized endogenous germ cell reporter mouse models, as well as a live ovarian culture system and high-resolution imaging technology, to develop a subcellular resolution 4D-whole ovary imaging system. Through this system, we were able to capture the entire process of oocyte selection in mouse ovaries and generate comprehensive visual data demonstrating an intense competition between oocytes, which is unrelated to cysts, to select the best oocyte candidates for long-term survival. These findings provide a holistic understanding of how female mammals establish a high-quality oocyte reserve over time to accommodate their reproductive lifespan.

## Results

### 4-dimensional imaging to record germ cell development in live mouse ovaries

To record the live behavior of germ cells during selection, we established a 4- dimensional (4D) organ imaging platform by combining an in vitro mouse ovarian developmental system with a 3D high-resolution live organ imaging system (Fig. 1a to c). In the system, we scanned fetal ovaries containing endogenous fluorescence-labeled germ cells under a low phototoxicity high-depth imaging system (Fig. 1a) and reconstituted 3D high-resolution imaging that contained detailed information about the morphology and spatial locations of germ cells in live ovaries (Fig. 1b). The ovaries were then cultured in vitro and regularly imaged to create 4D germ cell developmental dynamics covering the entire process of germ cell selection and follicle formation (Fig. 1c). This finally generated time-lapse images containing detailed information about behavior and developmental dynamics of germ cells during their fate determination in live ovaries (Movie S1).

**Figure 1.**
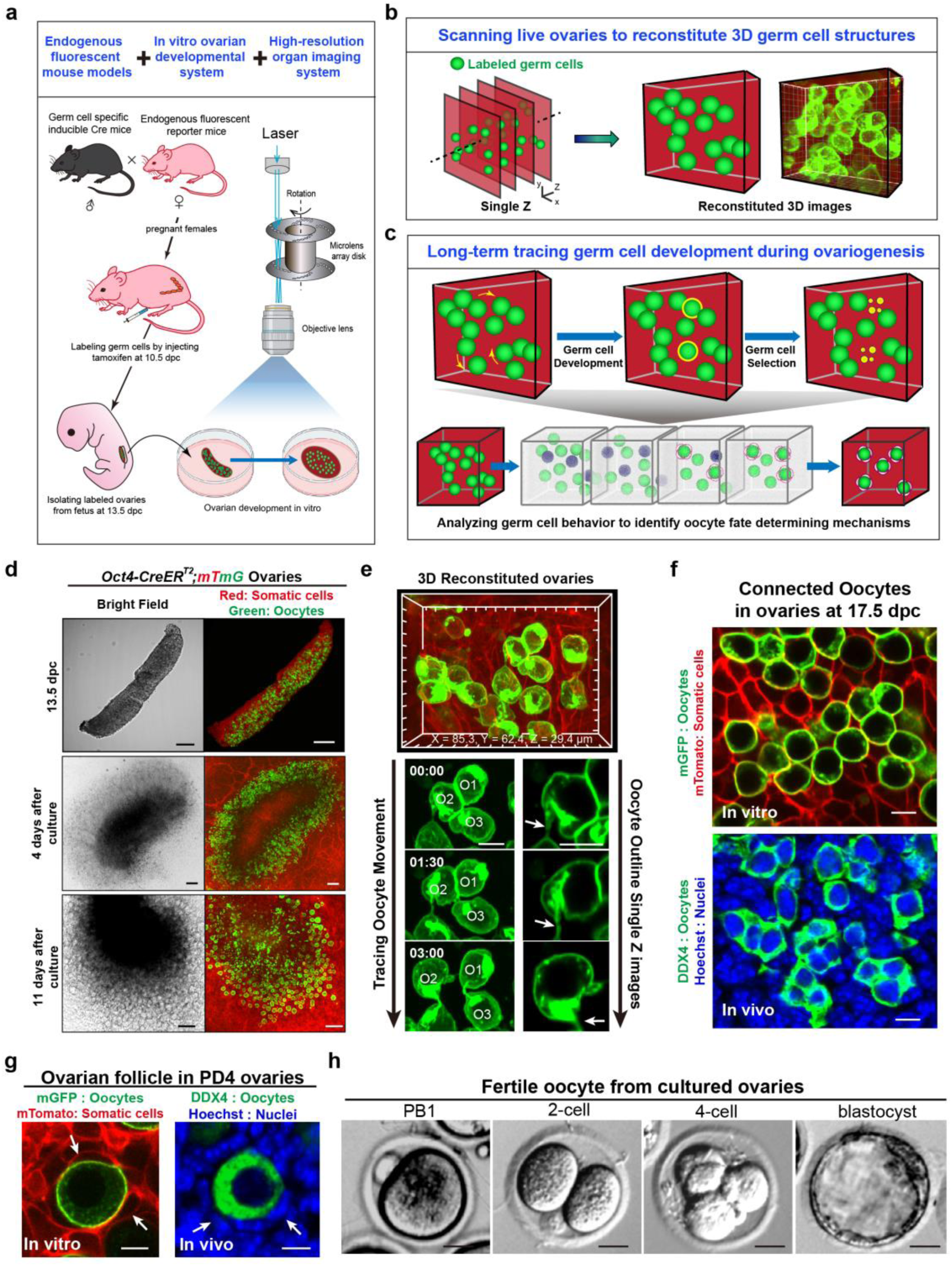
Establishing a 4D imaging system for recording germ cell development in live ovaries. **(a)** The schematic diagram illustrates the strategy for achieving high-resolution 4D imaging of germ cell development in fetal ovaries in vitro. The fetal ovaries which contained labeled germ cells were isolated and cultured in vitro, and then imaged under a high-resolution imaging system to record the developmental dynamics for a long-term tracing. **(b-c)** Schematic diagrams illustrate the reconstitution of 4D imaging of germ cell development in live ovaries. The ovaries were scanned to form Z-stack images with the detail information, which were then merged to reconstitute high- resolution, 3D ovarian images containing detailed information on germ cell morphology and spatial location (b). After long-term culture with continuous imaging, the 3D images at different time points were merged to form 4D developmental dynamics (c). **(d)** The morphology of in vitro developed ovaries at different periods. The ovaries were isolated from *Oct4-CreER^T2^;mTmG* females, in which the oocytes were labeled by mGFP (green) and other somatic cells were labeled by mTomato (red), at 13.5 dpc (top). After 4 days of culture, the ovaries were stabilized for imaging (middle), and large number of developed oocytes survived in ovaries after 11 days of culture (bottom). Scale bar: 100 μm. **(e)** The reconstituted 3D morphology of a group of oocytes (top). By tracing the developmental dynamics of these oocytes (bottom), the movement (left) and detail outline changes of these oocytes including subcellular structures (arrows) were recorded (right). Scale bar: 10 μm. **(f)** A single optical slide of cultured *Oct4-CreER^T2^;mTmG* ovaries at starting point of imaging, showing oocytes connected together forming cyst-like structures similar to those in fresh ovaries at 17.5 dpc. Scale bar: 10 μm. **(g)** At the ending point of imaging, oocyte covered by somatic granulosa cells (arrows) and formed follicle similar to that in fresh ovaries at PD 4. Scale bar: 10 μm. **(h)** In vitro fertilization of oocytes which were isolated from transplanted culture-ovaries. The oocytes were capable of forming healthy embryos at the 2-cell, 4-cell and blastocyst stages. Scale bar: 20 μm.

Since the major process of germ cell selection occurs between 17.5 dpc (day post coitum) to PD4 (Postnatal day) in mice ^23^, we designed a strategy to collect ovaries at 13.5 dpc (Fig. 1d, top) and cultured them for 4 days to stabilize the tissue (Fig. 1d, middle). Because of most germ cells have entered into meiosis after 17.5 dpc in the mouse ovaries ^23^, the germ cells were termed of oocytes in this study. The ovaries were then imaged for 7 consecutive days using the platform until follicle formation (Fig. 1d, bottom). To ensure the resolution of the oocytes, we first imaged the ovaries which isolated from an oocyte endogenous multi-fluorescent mouse model ^10,24^, *Oct4- CreER^T2^;mTmG* mice, in which the membrane of all oocytes is specifically labeled by membrane-GFP (mG) (Extended Data Fig. 1a-b) after tamoxifen treatment at 10.5 dpc. This allowed us to precisely identify the movement of oocytes (Fig. 1e, bottom left) and any changes in the outline of oocytes, including subcellular membrane structures (Fig. 1e, bottom right, arrows) in the 3D reconstituted ovaries (Fig. 1e).

Validating experiments showed that the system well-supported ovarian organogenesis and follicle formation in vitro. At the starting point of imaging (13.5 dpc + 4 days, equivalent to 17.5 dpc, and defined as c-17.5 dpc), we observed oocyte cyst structures (Fig. 1f, top) in the imaged ovaries, which were similar in morphology to oocytes in fresh ovaries at 17.5 dpc (Fig. 1f, bottom). At the endpoint of imaging (13.5 dpc + 11 days, equivalent to PD4 and defined as c-PD4), we observed ovarian follicles consisting of oocytes and surrounding (pre)granulosa cells (Fig. 1g, arrows), indicating the establishment of ovarian reserves in the cultured ovaries. Further allo-transplantation experiments showed that follicles in the in vitro-developed ovaries were healthy to growth (Extended Data Fig. 2) and able to ovulate fertilized oocytes (Fig. 1h), demonstrating that our system supported a normal process of oocyte development and ovarian reserve construction.

### Oocytes acted as single cells for their final fate determination

After validation, we traced oocyte development in *Oct4-CreER^T2^;mTmG* ovaries to analyze their detail behaviors during selection. Totally 1,099 EGFP-oocytes in six labeled ovaries were imaged from c-17.5 dpc and traced for 162 hours with 1.5 hours interval (Fig. 2a, Extended Data Fig. 3a and Movie S1). The time-lapse tracing successfully recorded the complete developmental dynamics of 594 oocytes, and statistical analysis revealed that the majority of the traced oocytes were eliminated (Fig. 2b, red), while 14.7 ± 2.7% of oocytes survived until c-PD4 (Fig. 2b, green) to form ovarian follicles. By analyzing the 4D dynamics of oocytes development, we recorded distinct developmental fates of different oocytes, including growth to survive (Fig. 2c, Lane 1, arrows), shrinking to elimination (Fig. 2c, Lane 2, arrows) and degradation to small pieces (Fig. 2c, Lane 3). Furthermore, subcellular changes of oocytes, such as oocyte deformation (Fig. 2c, Lane 4) and formation of oocyte filopodia (Fig. 2d, arrows), were also detectable in the images. Thus, the time-lapse images provided a comprehensive and visual database for analyzing oocyte fate determination in live ovaries.

**Figure 2.**
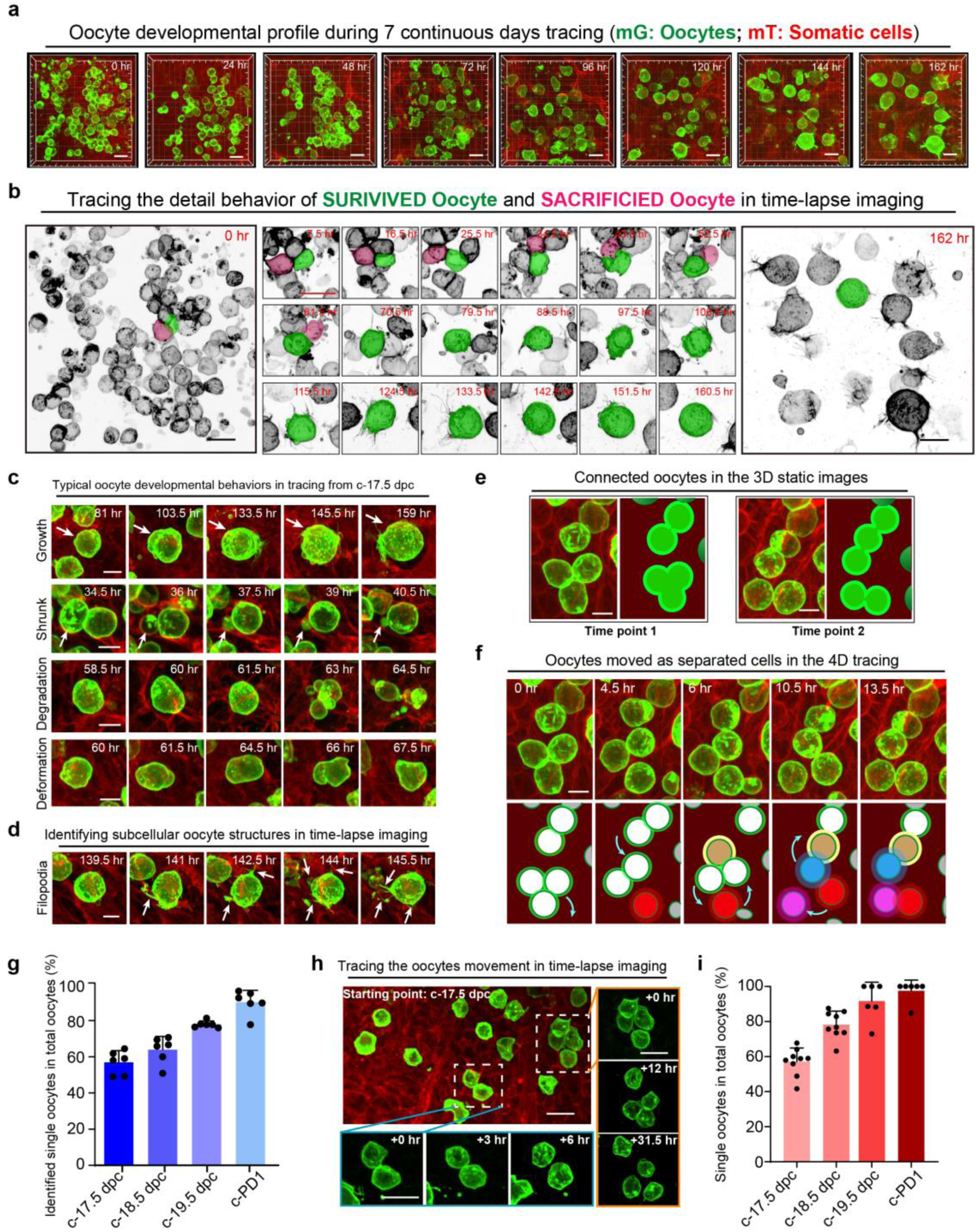
Oocytes act as separated single cells during their final fate determination. **(a)** A representative time-lapse image showing the oocyte developmental dynamics in live ovaries in vitro over a period of 162 hours, from c-17.5 dpc to c-PD4. Scale bar: 20 μm. **(b)** Tracing the developmental dynamics of oocytes in the time-lapse image. All oocytes were inverted to black/white (b/w), and a survival oocyte was colored green, while a sacrificed oocyte was colored red to highlight their developmental dynamics. Scale bar: 20 μm. **(c)** Representative images showing the typical behaviors of oocytes during development, including growth, shrinkage, degradation and deformation. Scale bar: 10 μm. **(d)** Representative images showing the filopodia formation of oocytes during development. Green, oocytes; Red, somatic cells. Scale bar: 10 μm. **(e)** 3D static images showing that oocytes connected to cyst-like structures at any detected time points. Green, oocyte; Red, somatic cells. Scale bar: 10 μm. **(f)** Tracing the oocyte movement in the 4D tracing images. Showing connected oocytes moved in different directions, and most of cyst-like structures were actually assembled by separated single oocytes in the ovaries after c-17.5 dpc. Scale bar: 10 μm. **(g)** Quantifying the ratio of separated oocytes in total oocytes from c-17.5 dpc to c-PD1. By continuously tracing the oocyte movement in the 4D time-lapse imaging, the ratio of oocytes with separation events in total oocytes was counted every 24 hours. Data are presented as the mean ± SD. **(h)** Identifying the movement of labeled oocytes in the *Oct4-CreER^T2^;mTmG* ovaries after low dosage of tamoxifen treatment. Only a small portion of cysts were labeled in the ovaries, with most of them breaking down into single oocytes at c-17.5 dpc. Green, oocytes; Red, somatic cells. Scale bar: 20 μm. **(i)** The ratio of single oocytes in total labeled oocytes from c-17.5 dpc to c-PD1 in ovaries with low labeling density of oocytes. Data are presented as the mean ± SD.

According to the classic model, oocyte fate determination is dependent on oocyte communication within the cyst ^2^. To investigate this, we first traced the development of oocytes within one cyst to identify any behavioral differences. Despite a high density of labeled oocytes crowding the ovaries (Fig. 2b), cysts with different numbers of connected oocytes (Fig. 2e) were still identifiable in 3D static images at all detected time points. However, unexpectedly, analysis of the 4D tracing revealed that the oocytes within the identified 3D cysts were not stably connected. As the ovaries developed, the oocytes were actively moving, and the attached oocytes in one cyst separated and moved in different directions to connect with other oocytes, forming a new group of cysts (Fig. 2f and Movie S2). These separation and re-connection events were frequently observed in cyst-like structures with the development of ovaries, suggesting that most of the cyst structures observed in the static images were temporary oocyte attachments from 17.5 dpc onwards (Fig. 2g).

To confirm our findings, we reduced the tamoxifen dosage and labeled a small portion of germ cells (18.5 ± 2.3%, Extended Data Fig. 4a-b) at 11.5 dpc to trace germ cell development in *Oct4-CreER^T2^;mTmG* mice. This strategy allowed us to trace the developmental trajectory of germ cells from distinguishable cysts (Extended Data Fig. 4a-b) with no disruption from surrounded cysts. Using the 4D imaging platform, we traced the movement of these labeled germ cells from c-17.5 dpc to c-PD1, and the tracing results clearly showed that most of the germ cells (57.1 ± 7.8%) were moving as single cells in the ovaries since c-17.5 dpc, and most of connected oocytes also separated in the followed 48 hours (Fig. 2h). Statistical analysis showed that the ratio of single oocytes increased from 57.1 ± 7.8% at c-17.5 dpc to 78.1 ± 7.7% at c-18.5 dpc and 91.6 ± 10.8% at c-19.5 dpc (Fig. 2i and Extended Data Fig. 4c). By c-PD1, almost oocytes (97.5 ± 6.2%) had separated into single cells, and few cysts with connected oocytes existed in the ovaries (Fig. 2i). This finding demonstrated that cyst breakdown completed much earlier than follicle formation in the mouse ovary and oocytes acted as independent cells rather than cysts in the last period of their fate determination. Hence, an uncovered cyst-independent system is necessary to complete the oocyte fate determination in the mouse ovaries.

### Robust competition occurred between oocytes to decide the survived winners

To identify the model of cyst-independent system for oocyte fate determination, we tracked the developmental dynamics of oocytes (Extended Data Fig. 5a) that formed ovarian follicles at the end of culture. Through time-lapse images, we discovered several key developmental characteristics of the cyst-independent oocyte selection process. Firstly, all the survived oocytes (referred to as winner-oocytes in this study) formed filopodia-like structures (FLs) on their cell surface during development (Fig. 3a, arrows, blue box). Secondly, multiple oocyte debris (ODs) with GFP membrane were observed surrounding the winner-oocytes, particularly during the period of FL formation (Fig. 3a, arrowheads, red box). By tracing the origin of ODs, we observed that some oocytes broke into small vesicles with a complete membrane (Fig. 3b, Oocyte with red color, Movie S3). These sacrificed oocytes were referred to as loser-oocytes, which play as the nurse cells in the oocyte fate determination. The counting results demonstrated that the formation of ODs peaked from c-19.5 to c-PD2 (Fig. 3c, red line), which corresponded to the peak period of winner-oocytes forming FLs (Fig. 3c, green line). These observations suggested an interested model of how single oocytes determine their fate, wherein intense competition occurs between separated oocytes and loser-oocytes sacrifice themselves to form various ODs for nursing winner-oocytes to enrich their cytoplasm for survival.

**Figure 3.**
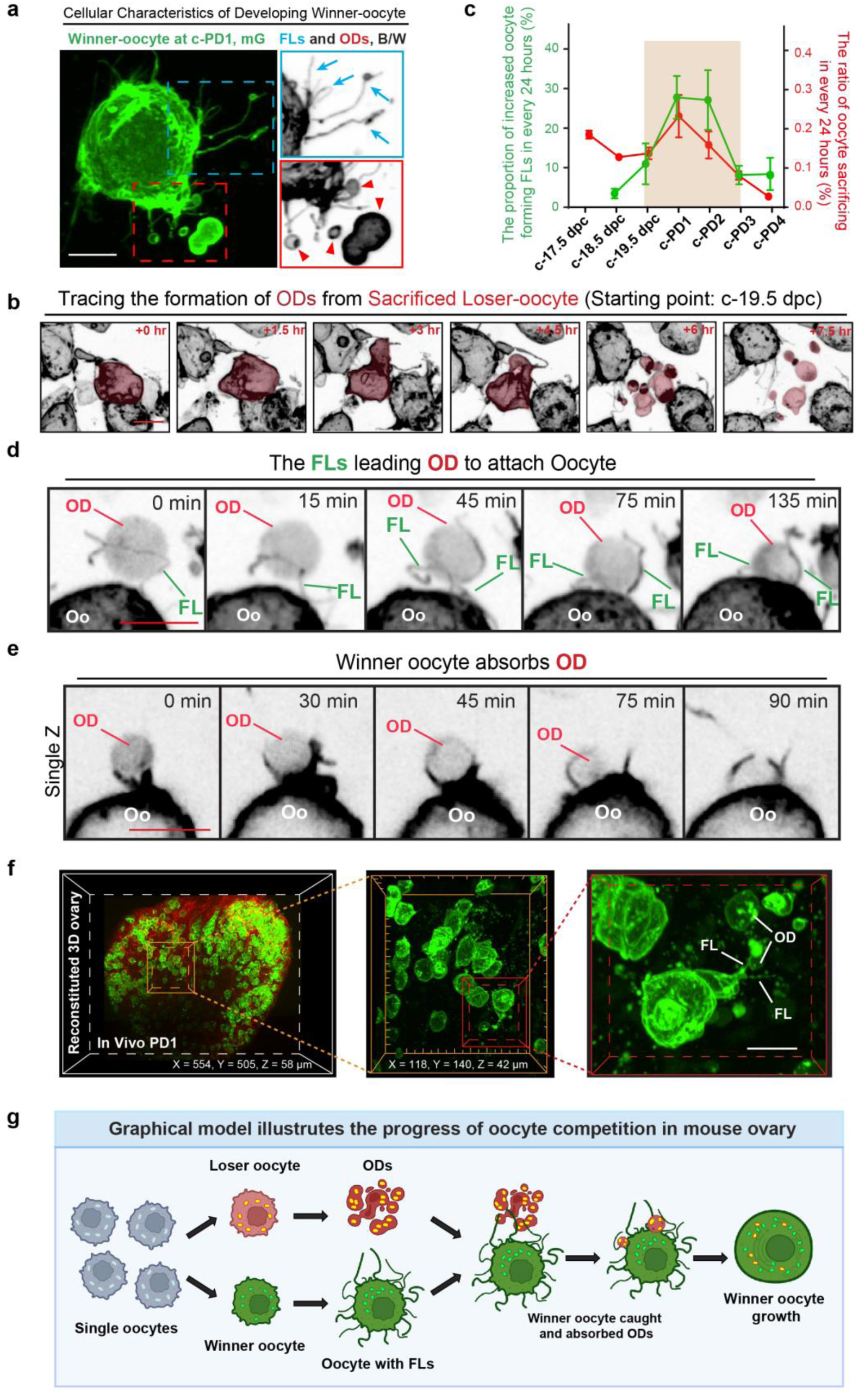
Intensive competition occurs between separated oocytes to decide their fate. **(a)** A representative image showing the developing cellular characteristics of survived oocytes (winner-oocyte). Highlighting the formation of filopodia-like structures (FLs, arrows) and surrounding by various oocyte debris (OD, arrowheads). Scale bar: 10 μm. **(b)** Tracing the derivation of ODs. The sacrificed oocyte (loser-oocyte) was colored red to highlight the progress of OD formation. Scale bar: 10 μm. **(c)** The frequency of winner-oocytes forming FLs (green line) and loser-oocytes forming ODs (red line), showing a consistent peaking period of two events (brown region) from c-19.5 dpc to c-PD2. Data are presented as the mean ± SEM. n = 4 ovaries. **(d)** The FLs from winner-oocytes extended towards the ODs to assist them in moving towards the winner- oocyte and then attached ODs. Scale bar: 10 μm. **(e)** Representative time-lapse images showing the detailed progress of how winner-oocytes absorbed the ODs. Scale bar: 10 μm. **(f)** FLs on the oocytes and ODs were observed in fresh transparent ovaries at PD1. Scale bar: 10 μm. **(g)** The graphic model illustrates the cyst-independent oocyte competition in mouse ovaries. The loser-oocyte sacrifices itself to form various ODs containing organelles enriched cytoplasm. The ODs are caught by winner-oocyte formed FLs to move forwarding to the winner-oocyte. Then the winner-oocyte absorbs the ODs to enrich cytoplasm and organelles for survival. Green, winner oocyte; Red, loser oocyte; Dark red, ODs.

To confirm our proposed model and to capture the detail oocyte behaviors during competition, we increased the frequency of photography to every 15 minutes for analyzing the oocyte development from c-PD1 over a 12-hour period (Movie S4). The resulting 4D tracing images revealed that winner-oocytes extended FLs towards surrounding ODs, directing ODs towards themselves (Fig. 3d, top and Extended Data Fig. 5b). Subsequently, winner-oocytes absorbed the ODs, facilitated by process structures in their membrane (Fig. 3e and Extended Data Fig. 5c), resulting in a noticeable increase in oocyte size due to cytoplasm enrichment (Extended Data Fig. 5d). These findings provide clear evidence that winner-oocytes “eat” the loser-oocyte formed ODs to ensure their survival.

Our findings have shown evidence of oocyte competition within cultured mouse ovaries. However, it remains uncertain whether this phenomenon is a naturally occurring physiological process. To confirm this point, we utilized a whole-mount ovarian transparent imaging system ^25^ to examine the presence of FLs and ODs in fresh *Oct4- CreER^T2^;mTmG* ovaries at PD1 (Fig. 3f left). Through 3D reconstituted ovarian images, we discovered that oocytes possess irregular cell shapes, much like those found in cultured ovaries. Significantly, we identified both FLs on oocytes and ODs (Fig. 3f, middle and right) in the ovaries at high resolution images. Moreover, we observed that some oocytes extended FLs in the direction of ODs, suggesting that FLs may have a role in catching the ODs (Fig. 3f, right). These evidences provide clearly confirmation that competition is a physiological event that occurs within mouse ovaries. Therefore, our results demonstrate that intense competition occurs between different single oocytes, and the winner ones appropriate the enriched organelles from ODs to enhance their quality for survival (Fig. 3g).

### Oocyte competition leaded cytoplasm exchanges nurse the winner-oocytes

Our finding demonstrated that winner-oocytes engulf ODs to enrich their cytoplasm, but it is unknown whether this is the primary strategy for cytoplasm exchange beyond cyst communication. To investigate this, we examined the strategy of oocyte cytoplasm exchanges by introducing an oocyte cytoplasm tracing mouse model *Oct4- CreER^T2^;Rainbow* mice into the system. In the ovaries of *Oct4-CreER^T2^;Rainbow* females (Extended Data Fig. 6a-b), the germ cells randomly expressed CFP or RFP in their cytoplasm (Fig. 4a), therefore cytoplasm exchange could be directly visualized through mixed fluorescence in cells (Fig. 4a, arrow). By monitoring the color of oocytes, we observed a small portion of oocytes with mixed fluorescent cytoplasm (purple) at all time points from c-17.5 to c-PD4 (Fig. 4b, arrows), and the ratio of purple oocytes showed that the peak of cytoplasm exchange occurred from c-PD1 to c-PD3 (Fig. 4c), following the peak of FLs and ODs formation. This finding suggested that oocyte competition is the primary, if not the only, approach to cytoplasm exchange between individual oocytes.

**Figure 4.**
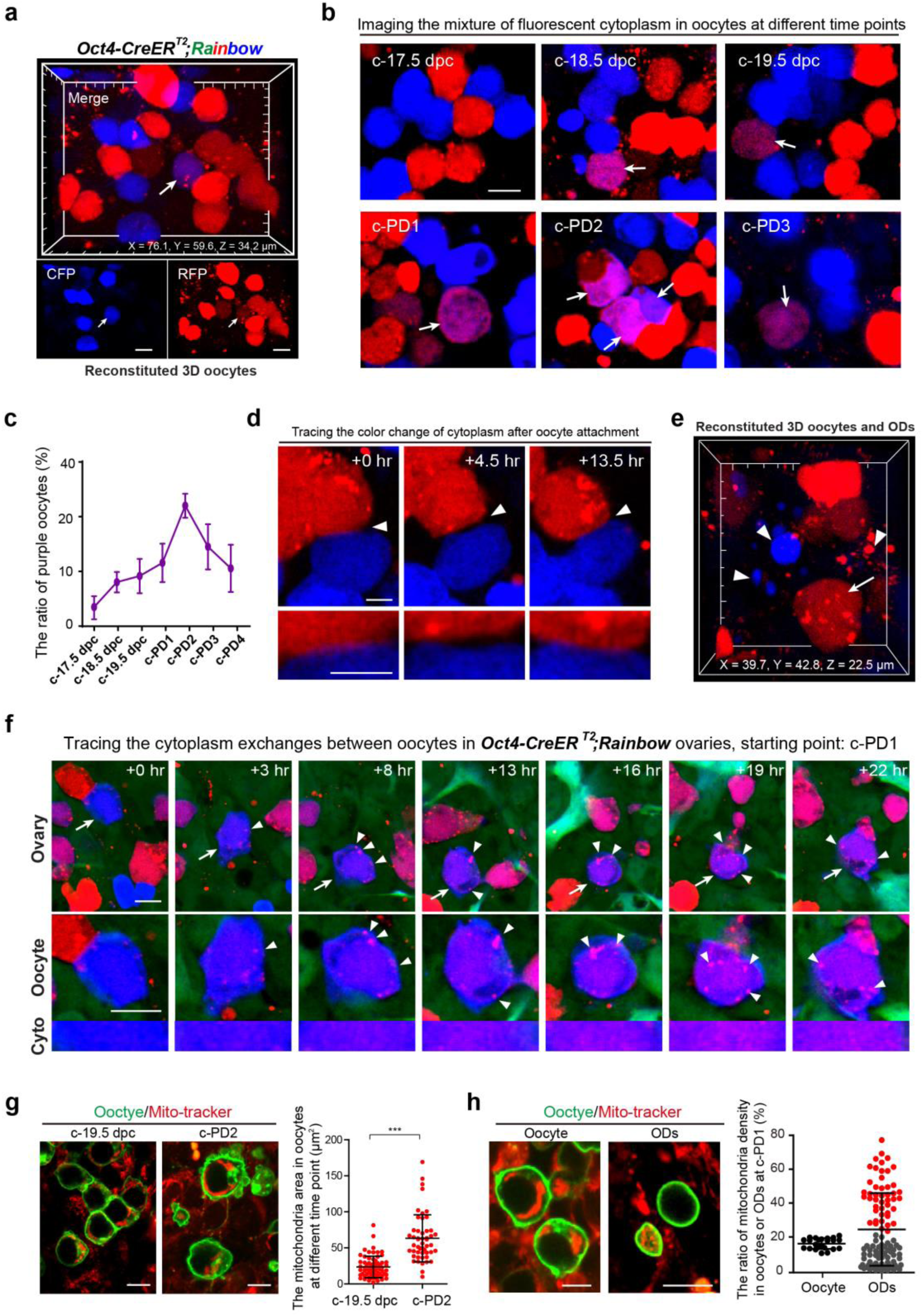
Oocyte competition is the major strategy for cytoplasm exchanges after cyst breakdown. **(a)** 3D reconstructed images of *Oct4-CreER^T2^;Rainbow* ovaries are shown, displaying random expressions of RFP (red) or CFP (blue) in oocyte cytoplasm. Cytoplasm exchange leads to a mixture of fluorescence in oocyte (purple, arrow). Scale bar: 10 μm. **(b)** Analysis of oocytes fluorescence reveals an increased number of oocytes with mixed fluorescent cytoplasm (purple, arrows) from c- 17.5 dpc to c-PD3. Scale bar: 10 μm. **(c)** The quantification of the ratio of purple oocytes in total oocytes from c-17.5 dpc to c-PD4, showing the peak of cytoplasm exchanges occurs between c- PD1 to c-PD3. Data are presented as the mean ± SD. **(d)** Tracing the attachment between oocytes with different fluorescence in the time-lapse imaging. No cytoplasm exchanges occurs after attachments (arrowheads) between oocytes. Scale bar: 5 μm. **(e)** The reconstructed 3D images demonstrate the existence of ODs (arrowheads) surrounding oocyte (arrow) in the *Oct4- CreER^T2^;Rainbow* ovaries. **(f)** The time-lapse images trace the progress of a blue winner-oocyte (arrow) absorbing red ODs (arrowheads), leading to a gradually change of cytoplasm color from blue to purple. The bottom line highlights the color change of winner-oocyte cytoplasm. Scale bar: 10 μm. **(g)** Analyzing the absolute area of mitochondria in oocytes (red: mitochondria, green: oocytes). Showing a significantly increased average mitochondria area in oocytes at c-PD2 compared to that in oocytes at c-19.5 dpc. Scale bar: 10 μm. Data are presented as the mean ± SD. ***P < 0.001, by two-tailed unpaired Student’s t-test. **(h)** Comparing the relative density of mitochondria in oocytes and ODs (red: mitochondria, green: oocytes or ODs) at c-PD1 (left). A great variation of mitochondria density was observed in ODs but not in oocytes (right). Scale bar: 10 μm. Oocyte: n = 23; ODs: n = 107. Data are presented as the mean ± SD.

In the classic model, oocyte connections are essential for cytoplasm exchanges^4^. Given high frequency of oocyte attachments and reassembly observed after cyst breakdown, we examined whether cytoplasm exchange occurred through connections with attached separated oocytes. By analyzing high magnification images, we traced a total of 100 attachments between oocytes with different colors (Fig. 4d, arrowheads), but did not observe any mixing of cytoplasm after the connections of oocytes (Movie S5). Furthermore, we observed small colorful ODs (Fig. 4e, arrowheads) that surrounded some oocytes (Fig. 4e, arrow), and 4D tracing analysis clearly captured that oocyte with single color fluorescent protein (Fig. 4f, blue oocyte, arrow) absorbed various ODs with another color fluorescence (Fig. 4f, red ODs, arrowheads and Movie S6) which led the progressive color change of oocyte from blue to purple (Fig. 4f, Cyto). This data clearly demonstrates that cytoplasm exchanges do not rely on connections and attachments between oocytes after the cyst breakdown, suggesting that oocyte competition is the only strategy for cytoplasm and organelles enrichment during the final period of oocyte selection.

Corroborating this observation, we identified mitochondrial in the oocytes in *Oct4- CreER^T2^;mTmG* ovaries and we noted that oocytes showed a marked increase in mitochondrial density from c-19.5 dpc to c-PD2 (Fig. 4g), which is the period of ODs and FLs formation peak. Furthermore, quantitative analysis revealed considerable variation in mitochondrial density among different ODs (Fig. 4h), suggesting that not only cytoplasm but also organelles, such as mitochondria, were enriched in loser-oocyte derived ODs and transferred to winner-oocytes.

### Single cell sequencing suggesting the competition was controlled by autophagy

To dissect underlying mechanisms that control oocyte competition, we performed single cell RNA-seq to identify molecular expressing characteristics in oocytes at PD1, the time of intensive competition occurred in ovaries (Fig. 5a). A total of 1,307 oocytes expressing an average of 4,084 genes were retained for analysis after stringent quality control. Then, the highly variable genes were selected to perform a principal-component analysis (PCA) and KNN clustering, and the oocytes were separated to 6 clusters (C0 to C5) based on their gene expressing characteristics (Fig. 5b). We found that meiotic related genes including *Dmc1* ^26^, *Meiob* ^27^ and *Spo11* ^28^ are significantly high expressions in the C0 and C1, suggesting these oocytes are in active meiotic progress and should be in early developmental state (Fig. 5c to d, red and Table S1). We therefore defined them as the candidate oocytes that are waiting to be selected. Cluster 2 and 3 are highly expressing *Sohlh1* ^29^, *Ooep* ^30^ and *Dppa3* ^31^, which are the specific genes of oocytes in formed follicles (Fig. 5c to d, green and Table S1), showing these cells were selected and should be the winner-oocytes for forming follicles in ovaries. Further analyses identified that *Diaph3* (Formin)^32^, *Epn2* (Epsin)^33^ and *Vil1* (Villin)^34,35^, which are genes in charge of the regulations of microvilli or filopodia formation (Fig. 5c to d, blue and Table S1), are significantly high expressions in the C1, C2 and C3. These findings are in consistent to the cell behaviors of winner-oocytes in our 4D imaging, confirming these oocytes are the winner-oocytes in ovaries. In cluster 4 and 5, oocyte death related genes including *Trp53* ^21^, *Anxa5* ^36^ and *Vdac1* ^37^ were highly expressions (Fig. 5c to d, purple and Table S1), implying they were loser-oocytes that would be sacrificed.

**Figure 5.**
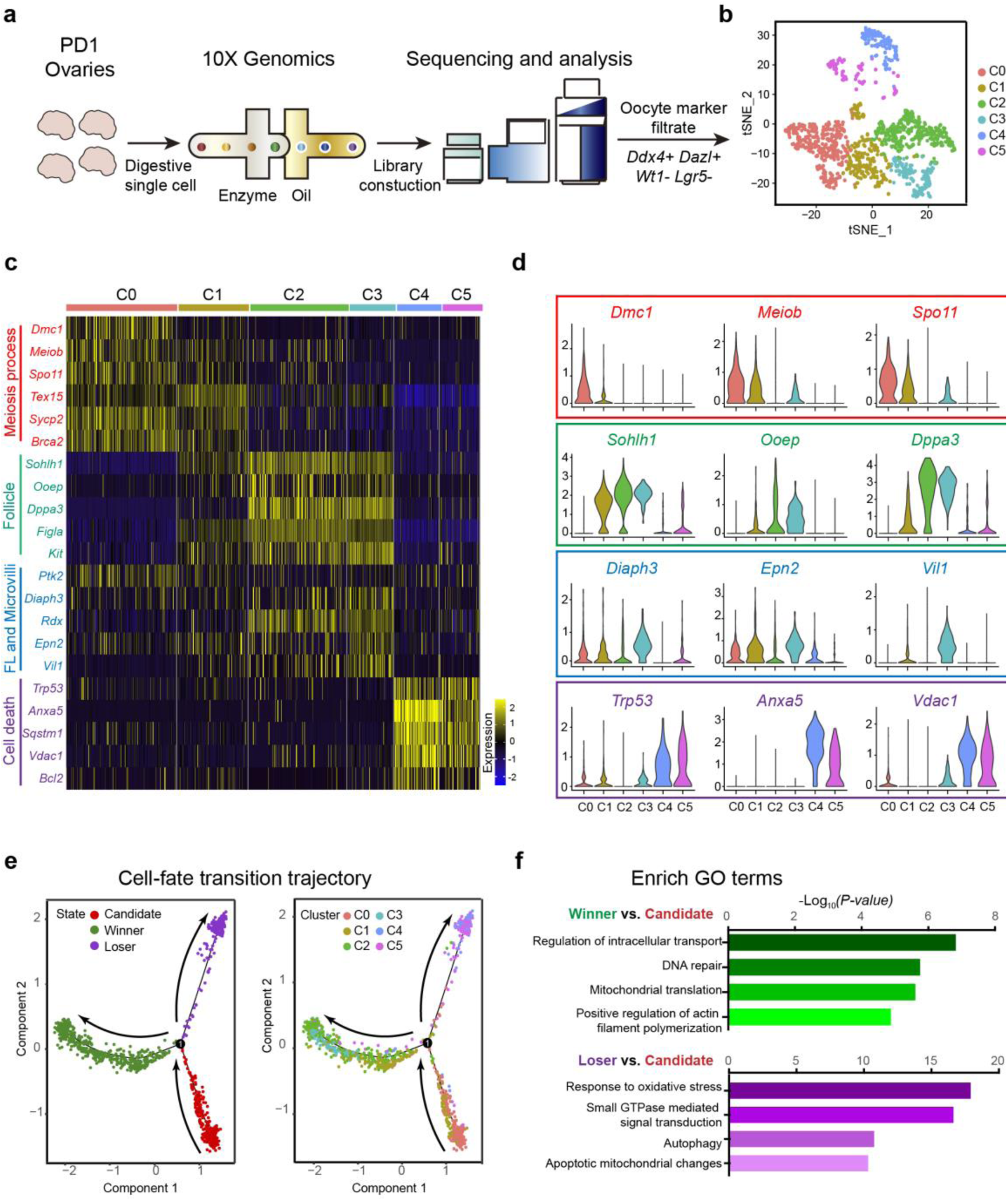
Single-cell RNA-Seq to analyze molecular mechanisms in regulating oocyte competition. **(a)** The flowchart provided an overview of the scRNA-seq for analyzing the gene expressing profile of oocytes in PD1 ovaries. **(b)** T-distributed stochastic neighborhood embedding (t-SNE) plots divided oocytes in PD1 ovaries into six separate clusters. **(c-d)** The gene expression signatures of various oocyte clusters were shown through a Heat map (c) and Violin plots (d). Meiosis process-related genes (red) gradually decrease from C0 to C5, with expression levels highest in the C0 and C1 clusters. Genes related to oocytes forming ovarian follicles (green) highly express in many oocytes in C2 to C3 clusters. FL and microvilli formation-related genes (blue) highly express in most oocytes across C0 to C3 clusters, with expression levels gradually rising from C0 to C3. Cell death-regulated genes (purple) highly express in most oocytes in C4 and C5. **(e)** Pseudotime prediction of the oocyte lineage trajectory, showing the cell lineage states (left), and cell cluster across the branches (right). The pseudotime scores were labeled by arrows. Cell lineage analysis and pseudotime dissection predicted the candidate oocyte lineage (red state in the left panel, C0 and C1 in the right panel) transitioned to the winner-oocyte lineage (green state in the left panel, C1, C2 and C3 in the right panel) and loser-oocyte lineage (purple state in the left panel, C4 and C5 in the right panel). **(f)** Representative gene ontology terms enriched in the winner-oocyte lineage (green histograms) and loser-oocyte lineage (purple histograms) are highlighted.

In consistent to gene expressing characteristics, lineage trajectory reconstruction according to the highly variable genes predicted that a clearly transcriptional map of oocyte fate determination (Fig. 5e), i.e. candidate oocyte population (C0 and C1, red state) differentiated to two distinguished cell lineages which were the winner-oocytes (C1, C2 and C3, green state) and loser-oocytes (C4 and C5, purple state). To further analyze how the lineages acquire their identities, we selected branch-related genes and analyzed the dynamics of gene expression along the predicted pseudotime from the candidate lineage to the winner-oocytes or loser-oocytes (Extended Data Fig. 7a). As shown in Extended Data Figure 7a to b, 399 genes were highly expressed in the candidate population, but their expression levels were dramatically reduced with the differentiation to the winner or loser populations (Extended Data Fig. 7a-b, red box and Table S2). With differentiation, the expression levels of 696 genes, including *Nobox* ^38^, *Uchl1* ^39^ and *Ybx2* ^40^ (Extended Data Fig. 7a-b, green box and Table S2), increased toward winner-oocyte lineage. The GO terms of these genes were enriched with “regulation of intracellular transport”, “DNA repair”, “mitochondrial translation” and “positive regulation of actin filament polymerization” (Fig. 5f, green bar-charts and Table S3). Meanwhile, we found that the expressions of 1,578 genes gradually increased with the differentiation of loser-oocyte lineage (Extended Data Fig. 7a-b, purple box and Table S4). These genes were enriched with GO terms on “response to oxidative stress”, “small GTPase mediated signal transduction”, “autophagy”, “apoptotic mitochondrial changes” (Fig. 5f, purple bar-charts and Table S4), suggesting that these genes or pathways might be crucial for the regulations of loser fate determination.

### Suppressing oocyte competition leaded a large but dis-functional ovarian reserve in ovaries

To figure out the physiological significance of oocyte competition in the ovaries, we chose to block autophagy which the genes were dramatic changed in loser-oocytes in our sc-seq analysis for disrupting the progress of oocyte competition. In the 4D imaging system, we added the well-tested autophagy inhibitor 3-MA (3-Methyladenine)^41^ from c-17.5 dpc, and traced the oocyte development and fate till c-PD4 (Fig. 6a). Generally, we found the oocytes in 3-MA group kept in a relative silence state with no active cellular development (Fig. 6a), growth (Fig. 6b) and death (Fig. 6c). Detail cellular analysis showed that the frequencies of both the behaviors of oocyte forming FLs (Fig. 6d, arrows and Fig. 6e) and formation of ODs (Fig. 6d, arrowheads) were dramatically decreased in the treated ovaries compared to those in controls. In consistent to the inactivity of oocyte competition, we found significantly increased number of oocytes at the end of imaging of c-PD4 in 3-MA treated ovaries (Fig. 6f, control 3,768 ± 709 vs. 3-MA 5,996 ± 386), and statistical analysis showed that both the average oocyte size (Extended Data Fig. 8a) and the density of mitochondria in survived oocytes (Extended Data Fig. 8b) were dramatically decreased in the treated ovaries. These results demonstrated that lacking of oocyte competition suppressed the selection of oocytes and might decline the oocyte quality of ovarian reserve.

**Figure 6.**
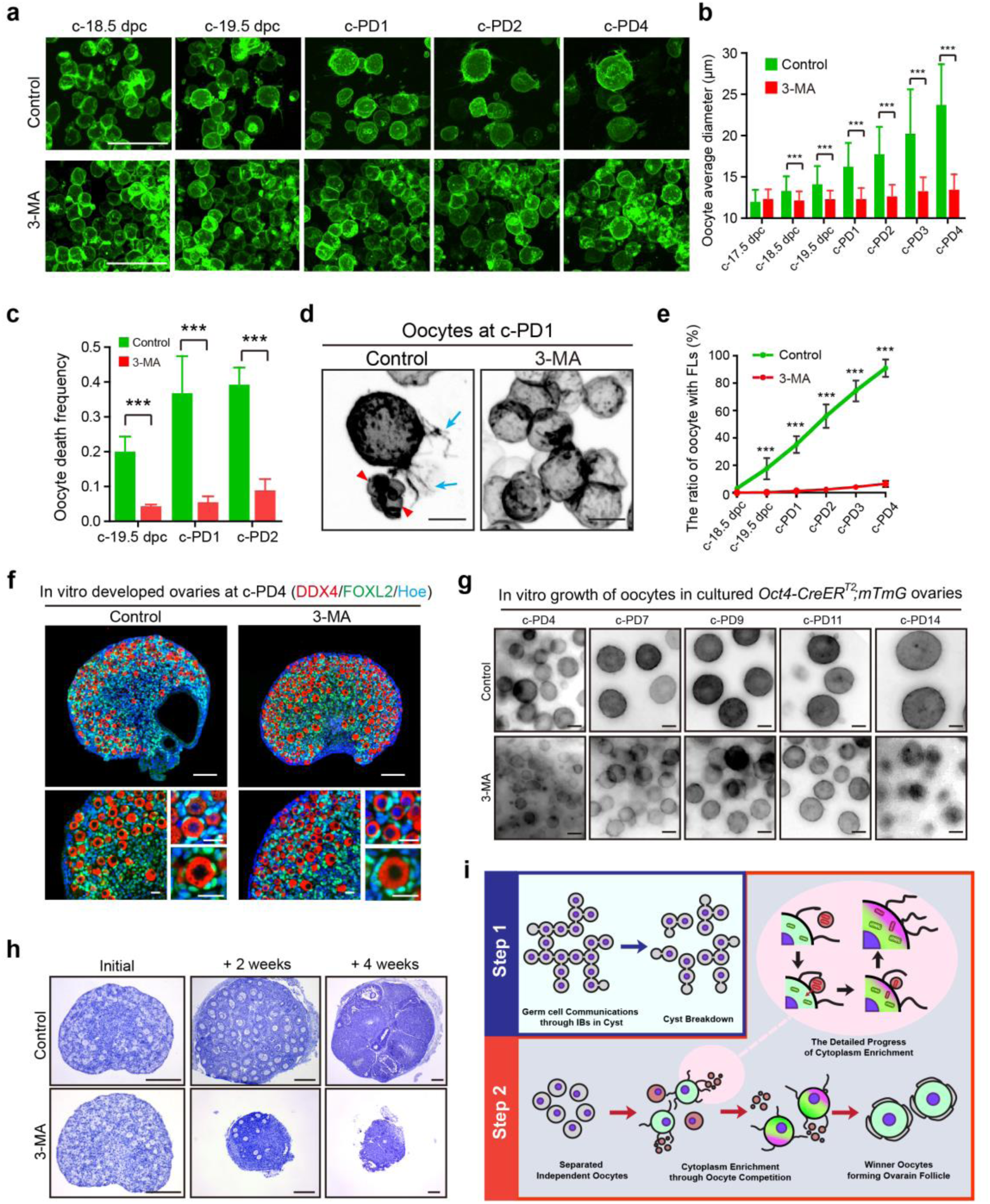
Blocking oocyte competition leads functional failure of ovarian reserve construction. **(a)** Tracing the oocyte development with or without 3-MA treatment. Normal oocyte competition and selection were observed in control ovaries, but not in the 3-MA treated ovaries. More oocytes with a dramatically decreased size survived at the end of culture at c-PD4. Scale bar: 50 μm. **(b)** Quantification of oocyte average diameter shows significant growth retardation in oocytes from 3- MA treated ovaries compared to those from control ovaries. More than 50 oocytes were measured for each time points. **(c)** Statistical analysis demonstrates a dramatically decreased frequency of oocyte death in 3-MA treated ovaries compared to that in control ovaries from c-19.5 dpc to c-PD2. n = 4 ovaries every group. **(d)** Representative images display the morphology of oocytes in 3-MA treated ovaries and control ovaries at c-PD1. Winner-oocytes with FLs (arrows) and loser-oocyte forming ODs (arrowheads) were observed in the control ovaries but not in 3-MA treated ovaries. Scale bar: 10 μm. **(e)** Statistical analysis of the oocytes with FLs shows that only a few oocytes formed FLs in 3-MA treated ovaries. More than 100 oocytes were measured for each time points. **(f)** Immunofluorescent staining displays ovarian morphology at c-PD4. More oocytes survived in 3-MA treated ovaries compared to that in control ovaries. Magnified images show ovarian follicles formed in both 3-MA group and control group. Red: DDX4, Green: FOXL2, Blue: Hoechst. Scale bar (top): 100 μm; Scale bar (bottom): 20 μm. **(g)** Investigation of the growth capability of oocytes derived from ovaries with or without 3-MA treatment. Although 3-MA was removed, oocytes in 3- MA treated ovaries were unable to fully grow. Scale bar: 10 μm. **(h)** Histological analysis of ovarian development after 3-MA treatment. Normal follicle development and corpus luteum formation were observed in control ovaries during 4 weeks of in vivo development. In the 3-MA group, only a few oocytes survived after 2 weeks of transplantation, and no healthy follicles were found in ovaries at 4 weeks after surgery. Scale bar: 200 μm. **(i)** The model of two-step oocyte selection in mammals. In the first round of cyst-dependent selection, orderly cytoplasm exchange occurs in cysts between intercellular bridges to improve the quality of selected germ cells. After cyst breakdown, the second round of selection which is cyst independent occurs through intense oocyte competition between individual oocytes. The winner-oocytes form FLs to capture organelles enriched ODs, which are derived from loser-oocytes, to enhance their quality. The best oocytes survive through two-step oocyte selection, constructing the ovarian reserve that supports female fertility throughout their life. The colors were inverted to black/white (b/w) in (d) and (g) to highlight oocyte morphology. Data are presented as the mean ± SD. ***P < 0.001, by two-tailed unpaired Student’s t-test.

Next, we extended the time of in vitro culture to monitor the developmental capability of oocytes which lacked the competition. After 7 days of additional culture (equal to PD 11), oocytes in control ovaries kept growth and the average diameter of them increased to 52.4 ± 4.8 μm (Fig. 6g and Extended Data Fig. 8c, control), showing our system well support the growth of oocytes in vitro. However, the lacking competition oocytes just increased to 37.4 ± 6.1 μm (Fig. 6g and Extended Data Fig. 8c, 3-MA) at the end of culture. This data demonstrated that the oocytes without competition were disable to fully grow. Meanwhile, we performed allo-transplantation to examine in vivo developmental capability of the oocytes with or without competition (Fig. 6h), and we found that a fast loss of oocytes occurred in 3-MA treated ovaries after 2 weeks of transplantation (Fig. 6h, 3-MA + 2 weeks), and no healthy oocytes survived in the treated ovaries after 4 weeks of surgery (Fig. 6h, 3-MA + 4 weeks). This is in sharp contrast to the healthy development and ovulation of oocytes (Fig. 6h, control) in the control group after transplantation. Therefore, we concluded that the oocyte competition is essential to boost oocyte quality for the maintenance of normal female fertility in mice.

In summary, our study suggests a two-step selection model to boost oocyte quality in mammals (Fig. 6i). After cyst formation in the fetal ovaries, the germ cells are initial selected through the IBs in cysts ^4,12^ (Fig. 6i, step 1). With the cyst breakdown, intensive oocyte competition occurs between separated independent oocytes to complete the secondary selection, and finally decides the number of winner-oocytes for forming ovarian reserve (Fig. 6i, step 2). This specific competition system might be crucial for mammals to fit their long reproductive lifespan.

## Discussion

Oocytes are a specialized cell population responsible for generating new life and ensuring the continuation of a species, and complex processes of oogenesis have been evolved to ensure that ovulated oocytes are of the highest quality ^1,42^. One commonly understood mechanism involves the transfer of cytoplasm and organelles from sister nurse oocytes to the selected oocytes, thereby enhancing their quality ^4^. The classic model of cytoplasm exchange suggests that this process occurs in a gentle and orderly manner, solely dependent on oocyte connections within the cyst. Since all germ cells in a cyst are derived from a single progenitor germ cell, it appears that germ cell selection operates like a “primogeniture system”, where the original germ cell with the most intercellular bridges (IBs) is deemed the ultimate winner and develops into an ovulated egg ^2,12,43^. This model has been well-identified in insects especially in *Drosophila*, and recently studies demonstrated that it also fit in mice, suggesting this cyst-dependent gentle communication model is conserved from invertebrate to vertebrate species including mammals ^4^.

Our current findings reveal a cyst-independent oocyte competition system that ultimately determines the fate of oocyte survival in mice. These findings also suggest that a two-step germ cell selection model is responsible for constructing ovarian reserves in mammals. During ovariogenesis, germline cysts are formed following rapid division of primordial germ cells in fetal ovaries, leading to the first round of selection through IB communications within the cyst. In this selection process, cytoplasm transfers from nurse cells to the chosen oocytes, and subsequently, the nurse cells shrink and die, resulting in a decrease in the number of oocytes prior to parturition in mice ^12^. In the perinatal ovaries, a second round of selection is initiated by a highly intensive and chaotic oocyte competition after the cysts break down into individual oocytes. Unlike the orderly and cyst-dependent selection process of the first round, this oocyte competition is much more intense and individual oocytes compete vigorously for survival. By utilizing time-lapse tracing, we propose that the origin of this competition is derived from a burst of loser oocytes, controlled by autophagy. This leads to the redistribution of organelles into the cytoplasm of loser oocytes and eventual formation of oocyte-derived debris (ODs). These ODs are then caught by winner oocytes which generate follicle-like structures (FLs), acting as detectors to help the winner-oocytes merge ODs into cells to complete the enrichment of cytoplasm and organelles. After the two rounds of selection, the quality and quantity of the ovarian reserve are established, which determines the reproductive resources available to support female fertility throughout their lifetimes.

Our study on oocyte competition reveals that cyst-dependent communication is not the only means of enriching cytoplasm for oocyte survival in mammals. Furthermore, when autophagy is suppressed and oocyte competition is blocked, we observed an enlarged but incompetent oocyte pool in the ovaries. This suggests that oocyte competition is essential for enhancing oocyte quality in mammals. Our findings are consistent with previous observations that female mice with *Tex14* mutations (which disrupt the stability of IBs) are fertile but have a significantly reduced ovarian reserve size. In this mouse model, it seems that the first round of selection which is rely on the IB communications is lacking, but the second round of oocyte competition which is cyst independent should be occurred. Therefore, a decreased but fertile ovarian reserve is established in the female ovaries. These findings suggest that oocyte competition, which is a specific mechanism for oocyte selection in mice, plays a pivotal role in determining oocyte quality ^13,14^.

Our research has provided a dynamic depiction of the early development and fate determination of mammalian oocytes, revealing a previously unknown mechanism of preferential survival through intense oocyte competition during the selection process. Our findings highlight the need for more in-depth investigation of the intricate communication networks involved in oocyte development and selection. By gaining a comprehensive understanding of these mechanisms, we can gain valuable insights into factors that impact oocyte quality and the maintenance of reproductive function in female mammals.

## Materials and methods

### Animals

The C57 and ICR mice were obtained from the Laboratory Animal Center of the Institute of Genetics (Beijing). The *Rosa26^rbw/+^* and *mTmG* mice (007576, Jackson Laboratory) were generated as previously reported ^44,45^, and *Rosa26^rbw/+^* mice as a gift from Dr. Kui Liu. The *Oct4-CreER^T2^* mice ^24^ were gifted from Dr. Zheng Ping. In addition, to obtain *Oct4-CreER^T2^* mice, we designed polymerase chain reaction (PCR) primers for Cre enzyme DNA sequence (forward 1: 5′- CCAAGGCAAGGGAGGTAGACAAG -3′, forward 2: 5′- GCTTTCTCCAACCGCAGGCTCTC -3′, reverse: 5′- GCCCTCACATTGCCAAAAGACGG-3′). *Oct4-CreER^T2^* mice were crossed with *mTmG* and *Rainbow* mice to generate *Oct4- CreER^T2^;mTmG* and *Oct4-CreER^T2^;Rainbow*. All mice were housed in mouse facilities under 16/8-h light/dark cycles at 26 °C and humidity 40–70% with access to chow and water at libitum. The animal experiments conformed to the guidelines and regulatory standards of the Institutional Animal Care and Use Committee of China Agricultural University, No. AW72012202-3-1.

### Tamoxifen (Tam) administration to label oocytes

To collect the *Oct4-CreER^T2^;mTmG* and *Oct4-CreER^T2^;Rainbow* fetal ovaries, *mTmG* or *Rainbow* homogenous females at 6–8 weeks were mated with adult *Oct4-CreER^T^* males overnight. The presence of vaginal plugs in the following morning is counted as 0.5 dpc and the day after partum is defined as PD1. Tamoxifen (Tam, 75648, Sigma-Aldrich) was resuspended in 95% (v/v) ethanol (100 mg/ml) and then diluted in corn oil (C8267, Sigma-Aldrich) to a final concentration of 20 mg/ml. To label the oocytes in the ovaries of *Oct4-CreER^T2^;mTmG* and *Oct4-CreER^T2^;Rainbow* females, Tam, at a high dosage of 50 mg·kg^−1^ BW or low dosage of 20 mg·kg^−1^ BW, was intraperitoneally injected to pregnant mice at 10.5 dpc.

### In vitro development of fetal ovaries

The fetal ovaries were obtained from *Oct4-CreER^T2^;mTmG* and *Oct4- CreER^T2^;Rainbow* mouse fetuses at 13.5 dpc, and the mesonephros were separated and removed in precooled PBS (10 mM, pH 7.4) under a stereomicroscope (Stemi 305, Zeiss) with sterile conditions. To observe the detailed behaviors and development of oocytes, the fetal ovaries were cultured on 35 mm glass bottom dish (D35-14-1-N, Cellvis) with the DMEM/F-12 (Dulbecco’s Modified Eagle Medium/Nutrient Mixture F-12, 1320033, Invitrogen) supplemented with 1% ITS (insulin-transferrin-sodium selenite medium, I3146, Invitrogen), 10% FBS (fetal bovine serum, GIBCO, Life Technologies) and penicillin-streptomycin (100 IU/ml; 15140122, Invitrogen). After 4 days of adherent culture, the fetal ovaries (c-17.5 dpc) were attached firmly to the bottom of dishes for live cell imaging. The culture media was half-changed every other day to maintain the development of ovaries. The culture ovaries were photographed in a living cell workstation (Okolab) at 37°C, 5% CO_2_ by an Andor Dragonfly 502 spinning-disc confocal microscope for 7 days.

### High-resolution 4D ovaries live-cell long term imaging

To observe the detailed behaviors and development of oocytes in ovaries during perinatal period, images were acquired by an Andor Dragonfly 502 spinning-disc confocal microscope equipped with a 40× 1.3 NA, a sCMOS camera (Andor Zyla 4.2), and 488-nm (mG) and 568-nm (mT) lines of the Andor ILE system with a spinning-disc confocal scan head (Andor Dragonfly 502)^46^. The ovaries were acquired through Z-step mode with the index. In detail, images were acquired with laser 488-nm around 15 to 25%, laser 568-nm around 15 to 20%, exposure time 100 to 200 ms, and Z-step 0.7 μm for 30 to 50 μm by Fusion 2.1 software (https://andor.oxinst.com/products/dragonfly#fusion). To record the developmental dynamics of oocytes, 10-12 fields (308.72 x 308.72 μm^2^/field) in 4-6 ovaries were acquired at 1.5 interval hours for 162 continuous hours (Fig. 2a-f, Extended Data Fig. 3, Fig. 6a) or at 15 interval min for 12 hours (Fig. 3d and Extended Data Fig. 5b). Then, the z-stack images were merged and analyzed by Imaris software (https://imaris.oxinst.com/) to reconstitute 4D live oocyte developmental images.

To obtain the subcellular structures on oocytes in the imaging, the Z-stack images of time-lapse imaging were projected for about 30 μm by Imaris or Image J software (http://rsbweb.nih.gov/ij/) to reconstitute 3D oocyte live images. The ≥ 2.5 μm filopodia structure on oocyte was defined as the FL structure. The ODs were identified from time-lapse imaging, which oocyte burst to at least 3 debris (the diameter of ODs from 1.2 μm to 14 μm).

### Histological analysis and Immunofluorescence staining

For histological analysis, the ovaries were fixed in 4% paraformaldehyde (PFA, Santa Cruz, 30525-89-4) for 8 hr at 4 °C, dehydrated in ethanol and xylene, embedded in paraffin, then sectioned serially at 8 μm with a microtome (RM2245, Leica), deparaffinized and rehydrated. To observe ovarian morphology, sections were stained with hematoxylin (Santa Cruz, sc-24973A).

For immunostaining, the ovarian sections were first treated by high temperature (95- 98°C) for 16 minutes in 0.01% sodium citrate buffer (pH 6.0) to retrieve antigen. Then, 10% donkey serum (017-000-121, Jackson ImmunoResearch) was used to block the sections for 60 minutes at room temperature. Ovarian sections were incubated with different primary anti-bodies overnight at 4°C. The primary antibodies were as follows: FOXL2 antibody (IMG-3228, goat, 1:300, Novus), DDX4 antibody (ab27591, mouse, 1:200, Abcam) and TOM20 antibody (#42406, Rabbit, 1:200, Cell Signaling Technology). After PBS buffer washed, the sections were incubated with Alexa Fluor 555- or 488-conjugated donkey secondary antibody (1:100, Life Technologies) for 90 minutes at room temperature. Nuclear was counterstained by Hoechst 33342 (B2261, 1:100, Sigma). Images were acquired on Nikon Eclipse Ti digital fluorescence microscope or Andor Dragonfly spinning-disc confocal microscope.

### Ovarian allo-transplantation

Ovaries allo-transplantation was performed according to previously described protocols^47^. Generally, the cultured ovaries were transplanted under the kidney capsules of bilaterally ovariectomized adult females (6 to 8 weeks). The recipient females were anesthetized with Avertin (300 mg/kg; T48402, Sigma-Aldrich) and the bilaterally ovaries were surgical removed. Then, the kidneys were exposed and the capsule were cut to form an approximately 1-mm wound for seeding the tissues. The cultured ovarian was gently implanted into kidney capsule through the wound for the further in vivo development. After 2 or 4 weeks of surgery, the transplanted ovaries were collected to detect the development of ovaries and ovarian follicle distribution.

To detect the developmental potent of oocytes from transplanted ovaries, the recipient females were intraperitoneal injected 5 IU pregnant mare serum gonadotropin (PMSG, Sansheng Biological Technology) followed by 5 IU human chorionic gonadotropin (hCG, Sansheng Biological Technology) to stimulate ovulation after 4 weeks of ovarian transplantation. After 10 hr of hCG treatment, the transplanted ovaries were collected and the cumulus oocyte complexes (COCs) were puncture obtained from Graff’s follicles, and then in vitro fertilization (IVF) were performed.

### In vitro fertilization assay

The cumulus oocyte complexes (COCs) were collected as described above, and digested with 0.01% hyaluronidase (H-3757, Sigma) to gain cumulus-free oocytes. In vitro fertilization assay was performed as previously described protocols ^48^. The cumulus-free oocytes in HTF medium (MR-70-D, Merk-Millipore) were co-incubated with capacitated epididymal sperm of wildtype male mice in an incubator with 37°C and 5% CO_2_. Zygotes were washed and transferred to KSOM medium (MR-020P-5F, Merk-Millipore) for culturing to later stages after incubation for 6 hr. The embryos at different stages were acquired by Hoffman Modulation Contrast microscope (OLYMPUS IX71).

### High-resolution imaging of the subcellular structure on oocytes

To detect the labeling efficiency of oocytes in the *Oct4-CreER^T2^;mTmG* ovaries after Tam treatment, female gonads were collected at 11.5 dpc (Extended Data Fig. 4a-b) or 13.5 dpc (Extended Data Fig. 1b and Extended Data Fig. 4a-B) after different dosage of Tam injection at 10.5 dpc. To observation FLs and ODs in vivo, the *Oct4- CreER^T2^;mTmG* ovaries were collected at PD1 (Fig. 3g-h) after 30 mg·kg−1 BW of Tam injection at 12.5 dpc. The gonads and ovaries were fixed in 4% paraformaldehyde for 8 hr at 4 °C and washed with PBS containing 0.2% Triton X-100 and 0.5% 1- thioglycerol in the dark for 24 hours at room temperature. Next, the ovaries were incubated in the clearing medium (40% N-methylacetamide (M26305, Sigma-Aldrich) supplemented with 86% Histodenz (D2158, Sigma-Aldrich), 0.1% Triton X-100 and 0.5% 1-thioglycerol (M1753, Sigma-Aldrich) in the dark at room temperature on a rotor for 72 hours as previously described protocols ^25,49^. The clear *Oct4-CreER^T2^;mTmG* fluorescent gonads and ovaries were acquired by Andor Dragonfly spinning-disc confocal microscope with a 40× 1.3 NA Z-step 0.8 μm for 250-μm working distance objective as previously described indexes. Images were acquired by Fusion 2.1 software (https://andor.oxinst.com/products/dragonfly#fusion). For showing the detail morphology of FLs and ODs in vivo, the magnification images (Fig. 3f, middle and right) were crop from the 3D reconstructed image of whole transparent ovary.

### Mitochondria detection in cultured ovaries

To label mitochondria in oocytes in live ovaries, the *Oct4-CreER^T2^;mTmG* ovaries at c-17.5 dpc were incubated by MitoTracker™ Orange CMTMRos (1:1000, M7510, Thermofisher scientific) at 37°C, 5% CO_2_. After mitochondria staining, the ovaries were washed by DMEM-F12 medium with 10% FBS and then imaged by an Andor Dragonfly spinning-disc confocal microscope as previously described indexes.

### Single-cell dissociation of ovarian tissue

The approach of single-cell dissociation was modified from previous published method^50^. The ovaries from C57 mice at PD1 were collected and cut into small pieces. Then the ovarian pieces (∼5 ovaries/tube) were treated with 0.125% Trypsin (25200056, Thermofisher scientific) and 5 IU/ml RNase-free DNase I (EN0523, Thermofisher scientific) at 37 °C for total 15 minutes (5 min × 3), then neutralized in DMEM-F12 medium (11320-033, Gibco) with 10% FBS (16000-044, Gibco). After digestion, the mixture was filtered through a 40-μm cell strainer (352340, BD Biosciences) to avoid any cell aggregations and then cells were centrifuged at 300 g for 5 min. Removed the supernatant and resuspended cell-precipitation with DMEM-F12 medium and 10% FBS. To separation somatic cells and oocytes as described protocol ^51^, the ovarian single-cell mixture in 35 mm dish was incubated at 37°C, 5% CO_2_ for 8 hr. The supernatant contained oocytes was harvested and centrifuged at 300 g for 5 min. Removed the supernatant and resuspended cell-precipitation with DMEM-F12 medium (1% BSA).

### RNA-sequencing and primary sequencing analysis

For single cell RNA-sequencing, oocytes at PD1 were harvested as described above. Then cell viability and concentration were measured by LUNA (LUNA-FL^TM^, Logos biosystems) after AO/PI Cell Viability Kit staining (F23001, Logos biosystems). The oocytes viability was 85.9% and cell concentration was 1330 cells/µl. Then the cells were processed following 10x Genomics protocol ^50^. In brief, single-cell gel bead-in-emulsions (GEMs) were generated by Chromium Controller instrument (10x Genomics). After GEM-reverse transcriptions (GEM-RTs), GEMs were collected and the cDNAs were amplified and cleaned up with the SPRIselect Reagent Kit (Beckman Coulter). The libraries were constructed using the Chromium Single-Cell 3′ Library Kit version 3.1 (10x Genomics, Dual Index). Sequencing libraries were generated by NEBNext® UltraTM RNA Library Prep Kit for Illumina® (NEB, USA) and sequenced on an Illumina Novaseq platform by the Beijing Novogene (Beijing, China).

### Single-cell RNA-seq data processing

Barcode processing and gene counting were performed by Cell Ranger (v5.0.1) count pipeline with default parameters. Briefly, reads after trimming were mapped to the mm10 mouse genome (https://10xgenomics.com/) and the genome annotation (gencode vM23) with STAR packaged in the Cell Ranger, and then aligned reads were filtered for identifying cell barcodes and unique molecular identifiers (UMIs). As a result, 6,093 barcodes with a median of 9,879 UMIs per cell were got. The valid cell barcodes and UMIs were used for identifying cell clusters.

### Cluster and DEG analysis in single cell sequencing analysis

The R package Seurat (v4.1.0)^52^ were used to identify cell clusters and markers. First, cells were filtered by three criteria: the number of detected genes was greater than 100 and less than 7,500. The percentage of mitochondrial genes was less than 15%. For subsetting oocytes, the counts of *Ddx4* and *Dazl* oocyte genes was greater than 0 and the counts of *Wt1* and *Lgr5* somatic cell genes were less than 2. After filter, 1,307 cells were retained. Then, data normalization and scale were performed in order using function NormalizeData and ScaleData. The UMI counts of each gene were divided by the total UMI counts for that cell, multiplied by 10,000, and after natural-log transformed, the normalized data were scaled and centered. Then scaled data were performed dimensionality reduction using function RunPCA based on principal component analysis (PCA). Using a permutation test method based on null distribution (function ElbowPlot), we selected top 20 PCA dims to cluster cells by function FindNeighbors and FindClusters based on shared nearest neighbor (SNN) algorithm. To visualization the cell clusters, function RunTSNE was performed. Then, for each cluster, genes with an average log_2_-transformed difference greater than 0.25, a *P* value less than 0.01, and expressed in at least 25% cells were identified as cluster markers using function FindAllMarkers based on Wilcoxon rank sum test. These cluster markers (Table S1) were compared with widely accepted markers to help to determine the cell identities of each cluster.

### Pseudotime analysis

The R package monocle (v2.18.0)^53^ were used to infer potential differentiation trajectory of ovary mesenchymal cells. Briefly, after dimensionality reduction based on DDRTree algorithm, high variable genes identified by function differentialGeneTest were used to define cells in a pseudotime trajectory. All cells were divided into three states and ordered in three branches based on reversed graph embedding (RGE). The branch mostly made up of C0 and C1 was designated as root state, and then branch-related genes (Table S2) among high variable genes were shown in the heatmap using function plot_genes_branched_heatmap. The expression dynamics through pseudotime of representative genes were shown in Extended Data Figure 5b using function plot_genes_branched_pseudotime.

### GO enrichment analysis

The R package clusterProfiler (v3.18.1)^54^ were used to perform GO enrichment analysis. Multiple hypothesis test correction method was set to “BH”, and the *P* value and *q* value cutoff were set to 0.01 and 0.05, respectively. The results of GO enrichment analysis were attached in Table S3-4 and representative GO terms were show in corresponding figures.

### Autophagy inhibitor 3-MA treatment experiments

In the 3-MA treatment experiments, after 4 days of adherent culture, the *Oct4- CreER^T2^;mTmG* ovaries at c-17.5 dpc were treated with or without 3-MA (3- Methyladenine, 5 mM, M9281, Sigma-Aldrich)^41^ for 7 days, which media was full- changed every other day. The culture ovaries were imaged by an Andor Dragonfly spinning-disc confocal microscope as previously described indexes. After 7 days 3-MA treatment, the ovaries were washed by DMEM-F12 medium, and continued cultured in DMEM/F-12 supplemented with 10% FBS and 1% ITS. The ovaries were acquired by Nikon Eclipse Ti digital fluorescence microscope every other day for 7 days.

### Statistical analysis

To quantify the percent of single cell in live ovaries, we counted the numbers of separated single cell in every 24 hours (from c-17.5 dpc to c-PD1) of total oocytes in high (Fig. 2g) or low (Fig. 2i) tamoxifen induced *Oct4-CreER^T2^;mTmG* ovaries. In details, the movement of oocytes were traced in the reconstituted images, and the separation of germ cells was carefully identified by analyzing the 3D images. Germ cells with any potential connections (Extended Data Fig. 4d) were excluded from the counting of single cells.

To analyze the correlation of the formation of FL and OD structure, we quantified the difference value of FL ratio (the numbers of oocyte with FLs /total oocytes) and the frequency of oocyte death events (the numbers of oocyte death events/total oocytes) in every 24 hours from c-17.5 dpc to c-PD4 (Fig. 3c).

To analyze the correlation of winner oocyte growth and competition events in Extended Data Fig. 5c, we counted the total numbers of FLs on the oocyte of 33 image frames during c-19.5 to c-PD2 (as shown in x-axial) and measured the ratio of diameter of each oocyte at c-PD2/c-19.5 dpc (as shown in y-axial). The fitted curve (y = -4E-05x^3^ + 0.0034x^2^ - 0.0431x + 1.3572) was generated by Excel.

To identify the distribution of mitochondria in oocytes and ODs, we measured the area of mitochondria in oocytes (Fig. 3e) and the ratio of the mitochondria area/each oocyte or OD area (Fig. 3f).

To quantify the mixture cytoplasm of oocytes, we counted the percent of the numbers of purple oocytes in total oocytes every 24 hours from c-17.5 to c-PD4 (Fig. 4c).

To assess the effects of 3-MA for oocyte competition, we detected a series state indexes of oocytes with or without 3-MA treatment, including the diameter of oocytes every 24 hours c-17.5 to c-PD4 (Fig. 6b), the frequency of oocyte death events (the numbers of oocyte death/total oocytes) in every day from c-19.5 dpc to c-PD2 (Fig. 6c) and the percent of oocyte with FLs/total oocytes from c-18.5 dpc to c-PD4 (Fig. 6e).

All experiments were repeated at least three times. Sample organism participants were randomly allocated into experimental groups. In the process of analyzing experimental results, there was no blind assignment of researchers. Data are presented as the mean ± SD or SEM of each result. Data were calculated by Student’s t-test, and were regarded statistically significant at P < 0.05. P is suggested as follows: * P < 0.05, ** P < 0.01, *** P < 0.001 and not significant (n.s.), P ≥ 0.05. Statistics and charts were gained by using Prism 5 (GraphPad Software, La Jolla).

## Compliance and ethics

The authors have declared that no conflict of interest exists.

## Acknowledgements

The authors are grateful to Yun Li, Shan Jiang, Dr. Lan Jiang (Beijing Institute of Genomics, Chinese Academy of Sciences) for scRNA-seq technology support, Dr. Fengchao Wang (National Institute of Biological Sciences, Beijing, China) for IVF technology support, Dr. Kui Liu (Department of Obstetrics and Gynecology, The University of Hong Kong, Hong Kong, China) for kindly sharing the *Rosa26^rbw/+^* mice and Dr. Ping Zheng (Yunnan Key Laboratory of Animal Reproduction, Kunming Institute of Zoology, Chinese Academy of Sciences, Kunming, China) for kindly sharing the *Oct4-CreER^T2^* mice.

This study was supported by the National Key Research and Development Program of China to H.Z. and Y.Z. (2022YFC2703800), the Key Program of National Natural Science Foundation of China to H.Z. (82230051), the National Natural Science Foundation of China to Y.Z. (82371664), the 2115 Talent Development Program of China Agricultural University to Y.Z. (1021-00109022) and The Innovative Project of State Key Laboratory of Animal Biotech Breeding to H.Z. (2023SKLAB1-6).

## Author contributions

Y.Z. and H.Z. designed the research; Y.Z., L.J.,Y.B., X.X., L.L., W.W. and K.G. performed the experiments; Y.Z., Y.B., L.M., K.C., L.L., G.W., K.G., X.Y., L.J., X.X., Y.K., W.C., G.X. and H.Z. analyzed the data; Y.Z. and H.Z. wrote the paper. All authors have seen and approved the final version.

## Data availability statement

The raw sequence data reported in this paper have been deposited in the Genome Sequence Archive in National Genomics Data Center (GSA: CRA012740) that are publicly accessible at https://ngdc.cncb.ac.cn/gsa. The scRNA-seq data analysis was performed using scripts in R (version 4.1.0).

## Reference

1 Eppig, J. J. Oocyte control of ovarian follicular development and function in mammals. Reproduction 122, 829–838 (2001).

2 Lei, L. & Spradling, A. C. Mouse oocytes differentiate through organelle enrichment from sister cyst germ cells. Science 352, 95–99 (2016).

3 De Cuevas, M., Lilly, M. A. & Spradling, A. Germline cyst formation in Drosophila. Annual review of genetics 31, 405–428 (1997).

4 Spradling, A. C., Niu, W., Yin, Q., Pathak, M. & Maurya, B. Conservation of oocyte development in germline cysts from Drosophila to mouse. Elife 11, e83230 (2022).

5 Pepling, M. E. & C. Spradling, A. Female mouse germ cells form synchronously dividing cysts. Development 125, 3323–3328 (1998).

6 Cox, R. T. & Spradling, A. C. A Balbiani body and the fusome mediate mitochondrial inheritance during Drosophila oogenesis. Development 130, 1579–1590 (2003).

7 Bolívar, J. et al. Centrosome migration into the Drosophila oocyte is independent of BicD and egl, and of the organisation of the microtubule cytoskeleton. Development 128, 1889–1897 (2001).

8 Lei, L. & Spradling, A. C. Mouse primordial germ cells produce cysts that partially fragment prior to meiosis. Development 140, 2075–2081 (2013).

9 Findlay, J. K., Hutt, K. J., Hickey, M. & Anderson, R. A. How is the number of primordial follicles in the ovarian reserve established? Biology of reproduction 93, 111, 111–117 (2015).

10 Zhang, H. et al. Life-long in vivo cell-lineage tracing shows that no oogenesis originates from putative germline stem cells in adult mice. Proceedings of the National Academy of Sciences 111, 17983–17988 (2014).

11 Ikami, K. et al. Branched germline cysts and female-specific cyst fragmentation facilitate oocyte determination in mice. Proceedings of the National Academy of Sciences 120, e2219683120 (2023).

12 Niu, W. & Spradling, A. C. Mouse oocytes develop in cysts with the help of nurse cells. Cell 185, 2576–2590. e2512 (2022).

13 Greenbaum, M. P., Iwamori, N., Agno, J. E. & Matzuk, M. M. Mouse TEX14 is required for embryonic germ cell intercellular bridges but not female fertility. Biology of reproduction 80, 449–457 (2009).

14 Ikami, K. et al. Altered germline cyst formation and oogenesis in Tex14 mutant mice. Biology open 10, bio058807 (2021).

15 Baker, N. E. Emerging mechanisms of cell competition. Nature Reviews Genetics 21, 683–697 (2020).

16 Kim, W. & Jain, R. Picking winners and losers: cell competition in tissue development and homeostasis. Trends in Genetics 36, 490–498 (2020).

17 Li, W. & Baker, N. E. Engulfment is required for cell competition. Cell 129, 1215–1225 (2007).

18 Zhu, Y. et al. Migratory neural crest cells phagocytose dead cells in the developing nervous system. Cell 179, 74–89. e10 (2019).

19 Nagata, R., Nakamura, M., Sanaki, Y. & Igaki, T. Cell competition is driven by autophagy. Developmental cell 51, 99–112. e114 (2019).

20 Bowling, S. et al. P53 and mTOR signalling determine fitness selection through cell competition during early mouse embryonic development. Nature Communications 9, 1763 (2018).

21 Cochrane, C. R. et al. Trp53 and Rb1 regulate autophagy and ligand-dependent Hedgehog signaling. The Journal of Clinical Investigation 130, 4006–4018 (2020).

22 Ghafari, F., Pelengaris, S., Walters, E. & Hartshorne, G. M. Influence of p53 and genetic background on prenatal oogenesis and oocyte attrition in mice. Human Reproduction 24, 1460–1472 (2009).

23 Pepling, M. E. From primordial germ cell to primordial follicle: mammalian female germ cell development. Genesis 44, 622–632 (2006).

24 Greder, L. V. et al. Brief report: analysis of endogenous Oct4 activation during induced pluripotent stem cell reprogramming using an inducible Oct4 lineage label. Stem cells 30, 2596–2601 (2012).

25 Xu, X. et al. Imaging and tracing the pattern of adult ovarian angiogenesis implies a strategy against female reproductive aging. Science advances 8, eabi8683 (2022).

26 Bishop, D. K. RecA homologs Dmc1 and Rad51 interact to form multiple nuclear complexes prior to meiotic chromosome synapsis. Cell 79, 1081–1092 (1994).

27 Luo, M. et al. MEIOB exhibits single-stranded DNA-binding and exonuclease activities and is essential for meiotic recombination. Nature communications 4, 2788 (2013).

28 Romanienko, P. J. & Camerini-Otero, R. D. The mouse Spo11 gene is required for meiotic chromosome synapsis. Molecular cell 6, 975–987 (2000).

29 Pangas, S. A. et al. Oogenesis requires germ cell-specific transcriptional regulators Sohlh1 and Lhx8. Proceedings of the National Academy of Sciences 103, 8090–8095 (2006).

30 Pierre, A. et al. Atypical structure and phylogenomic evolution of the new eutherian oocyte- and embryo-expressed KHDC1/DPPA5/ECAT1/OOEP gene family. Genomics 90, 583–594 (2007).

31 Bortvin, A., Goodheart, M., Liao, M. & Page, D. C. Dppa3/Pgc7/stella is a maternal factor and is not required for germ cell specification in mice. BMC developmental biology 4, 1–5 (2004).

32 Schirenbeck, A., Bretschneider, T., Arasada, R., Schleicher, M. & Faix, J. The Diaphanous- related formin dDia2 is required for the formation and maintenance of filopodia. Nature cell biology 7, 619–625 (2005).

33 Aguilar, R. C. et al. Epsin N-terminal homology domains perform an essential function regulating Cdc42 through binding Cdc42 GTPase-activating proteins. Proceedings of the National Academy of Sciences 103, 4116–4121 (2006).

34 Friederich, E., Huet, C., Arpin, M. & Louvard, D. Villin induces microvilli growth and actin redistribution in transfected fibroblasts. Cell 59, 461–475 (1989).

35 Huelsmann, S., Ylänne, J. & Brown, N. H. Filopodia-like actin cables position nuclei in association with perinuclear actin in Drosophila nurse cells. Developmental cell 26, 604–615 (2013).

36 Ghislat, G., Aguado, C. & Knecht, E. Annexin A5 stimulates autophagy and inhibits endocytosis. Journal of Cell Science 125, 92–107 (2012).

37 Vijayan, M. et al. A partial reduction of VDAC1 enhances mitophagy, autophagy, synaptic activities in a transgenic Tau mouse model. Aging Cell 21, e13663 (2022).

38 Rajkovic, A., Pangas, S. A., Ballow, D., Suzumori, N. & Matzuk, M. M. NOBOX deficiency disrupts early folliculogenesis and oocyte-specific gene expression. Science 305, 1157–1159 (2004).

39 Goto, Y. et al. UCHL1 provides diagnostic and antimetastatic strategies due to its deubiquitinating effect on HIF-1α. Nature communications 6, 6153 (2015).

40 Zhang, Z. et al. YBX2-dependent stabilization of oocyte mRNA through a reversible sponge- like cortical partition. Cell Research, 1–4 (2023).

41 Wu, Y. et al. Synthesis and screening of 3-MA derivatives for autophagy inhibitors. Autophagy 9, 595–603 (2013).

42 Hsueh, A. J., Kawamura, K., Cheng, Y. & Fauser, B. C. Intraovarian control of early folliculogenesis. Endocrine Reviews 36, 1–24 (2015).

43 Soygur, B. et al. Intercellular bridges coordinate the transition from pluripotency to meiosis in mouse fetal oocytes. Science advances 7, eabc6747 (2021).

44 Zheng, W. et al. Two classes of ovarian primordial follicles exhibit distinct developmental dynamics and physiological functions. Human Molecular Genetics 23, 920–928 (2014).

45 Zhang, H. et al. Experimental evidence showing that no mitotically active female germline progenitors exist in postnatal mouse ovaries. Proc Natl Acad Sci U S A 109, 12580–12585 (2012).

46 Zhang, Y. et al. Oocyte-derived microvilli control female fertility by optimizing ovarian follicle selection in mice. Nature Communications 12, 2523 (2021).

47 Zhang, J. et al. In vivo and in vitro activation of dormant primordial follicles by EGF treatment in mouse and human. Clinical and Translational Medicine 10, e182 (2020).

48 Zhang, H. et al. Male fertility in Mus musculus requires the activity of TRYX5 in sperm migration into the oviduct. Journal of Cellular Physiology 235, 6058–6072 (2020).

49 Li, W., Germain, R. N. & Gerner, M. Y. Multiplex, quantitative cellular analysis in large tissue volumes with clearing-enhanced 3D microscopy (C(e)3D). Proc Natl Acad Sci U S A 114, E7321–e7330 (2017).

50 Mostovoy, Y. et al. A hybrid approach for de novo human genome sequence assembly and phasing. Nature methods 13, 587–590 (2016).

51 Teng, Z. et al. S100A8, An Oocyte-Specific Chemokine, Directs the Migration of Ovarian Somatic Cells During Mouse Primordial Follicle Assembly. Journal of Cellular Physiology 230, 2998–3008 (2015).

52 Satija, R., Farrell, J. A., Gennert, D., Schier, A. F. & Regev, A. Spatial reconstruction of single-cell gene expression data. Nature biotechnology 33, 495–502 (2015).

53 Qiu, X. et al. Reversed graph embedding resolves complex single-cell trajectories. Nature methods 14, 979–982 (2017).

54 Yu, G., Wang, L. G., Han, Y. & He, Q. Y. clusterProfiler: an R package for comparing biological themes among gene clusters. Omics : a journal of integrative biology 16, 284–287 (2012).

