## supplemental file for "Intense oocyte competition builds the female reproductive reserve in mice"

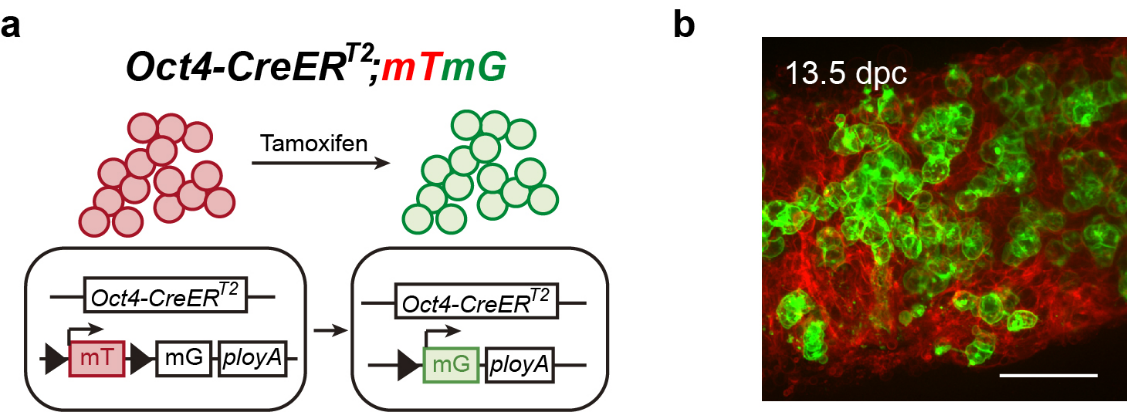

**Extended Data Figure 1. Labeling germ cell membrane in the *Oct4-CreER^T2^;mTmG* ovaries.**

**(a)** Illustration of tamoxifen (Tam)–induced labeling of germ cells in *Oct4-CreER^T2^;mTmG* ovaries. In *Oct4*-expressing germ cells, the CreER^T2^ recombinase is not active and the cells express mT, a membrane localized red fluorescent protein. Upon tamoxifen injection, the CreER^T2^ recombinase deletes the mT region and switches on the expression of mG, a membrane localized green fluorescent protein. Thus, the membrane of *Oct4*-expressing germ cell was labeled with green fluorescence. **(b)** The pregnant females with *Oct4-CreER^T2^;mTmG* fetus were given i.p. injection of 50 mg.kg^−1^ BW tamoxifen at 10.5 dpc, and the labeled fetal ovaries were collected at 13.5 dpc. Scale bar: 50 μm.

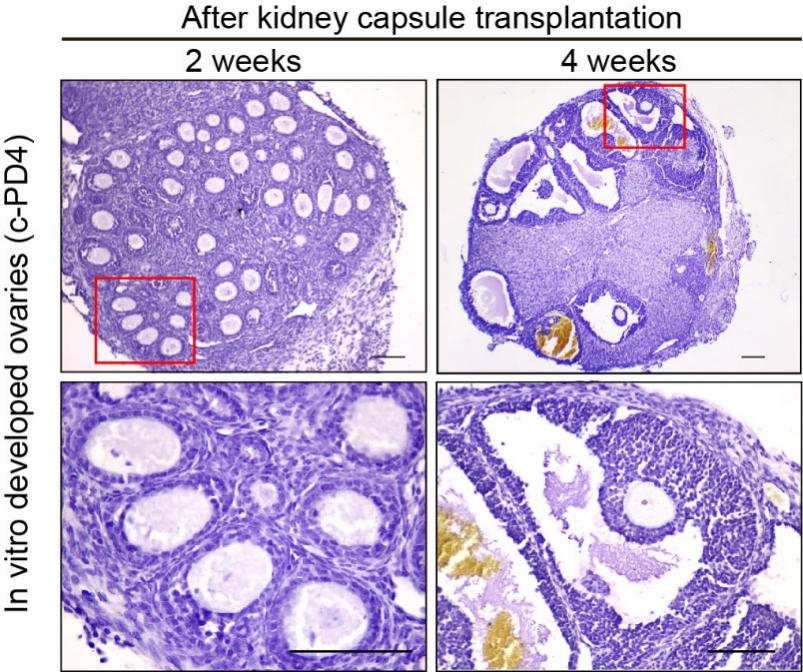

**Extended Data Figure 2. Normal follicle development in cultured ovaries after allo-transplantation.**

After in vitro culture, the ovaries were transplanted under the kidney capsule, and histological analysis showed that ovarian follicles developed normally after 2 and 4 weeks of transplantation. Scale bar: 100 μm.

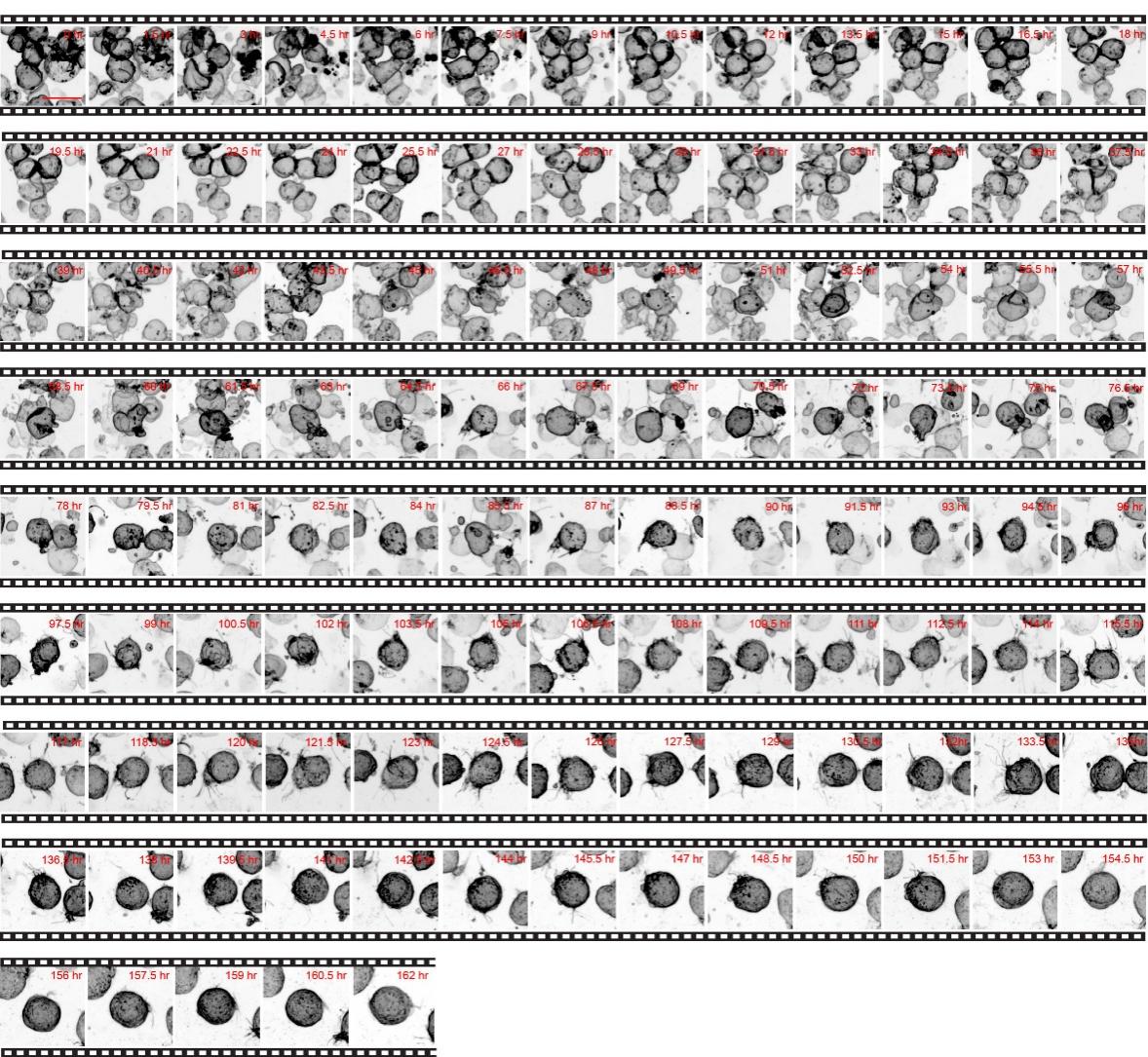

**Extended Data Figure 3. Time-lapse imaging to trace the development of a group of oocytes.**

The developmental dynamics of oocytes were traced in the time-lapse image with 1.5 interval hours for 162 hours. Oocytes were inverted to black/white (b/w). Scale bar: 20 μm.

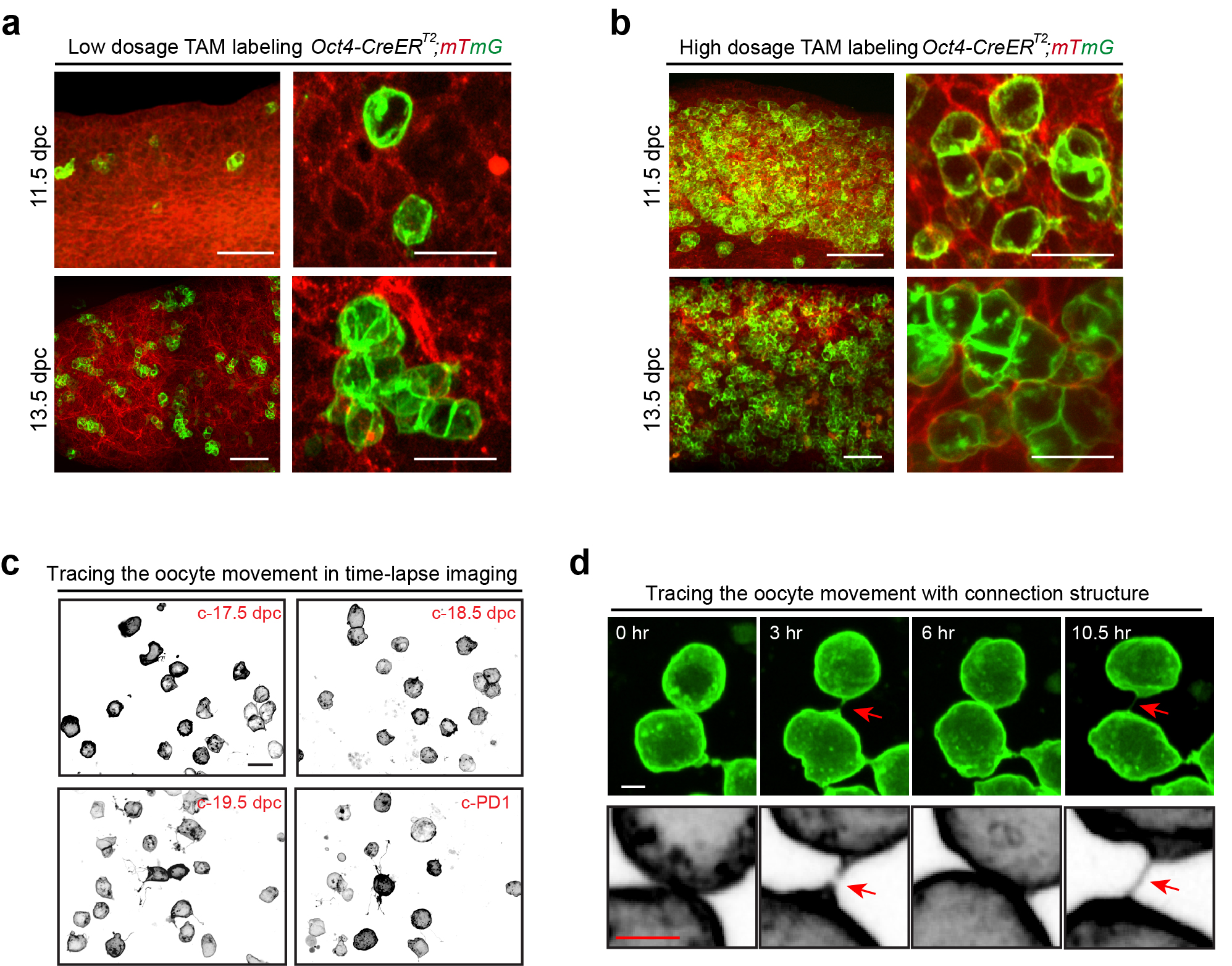

**Extended Data Figure 4. Tracing the oocyte movement in the *Oct4-CreER^T2^;mTmG* ovaries after low dosage Tam treatment.**

**(a-b)** Showing the labeling efficiency of oocyte in the *Oct4-CreER^T2^;mTmG* ovaries. The single germ cell at 11.5 dpc and separated cysts at 13.5 dpc were labeled in ovaries after low dosage of Tam treatment (a), whereas connected germ cells at 11.5 dpc and crowded cysts were observed at 13.5 dpc after high dosage of tamoxifen treatment (b). Scale bar: 50 μm. **(c)** Tracing the movement of labeled oocytes demonstrated majority of oocytes moving as single cell from c-17.5 dpc to c-PD1. Oocytes were inverted to black/white (b/w). Scale bar: 20 μm. **(d)** Germ cell movement with connections (arrows) were excluded from the counting of single cells. Scale bar: 5 μm.

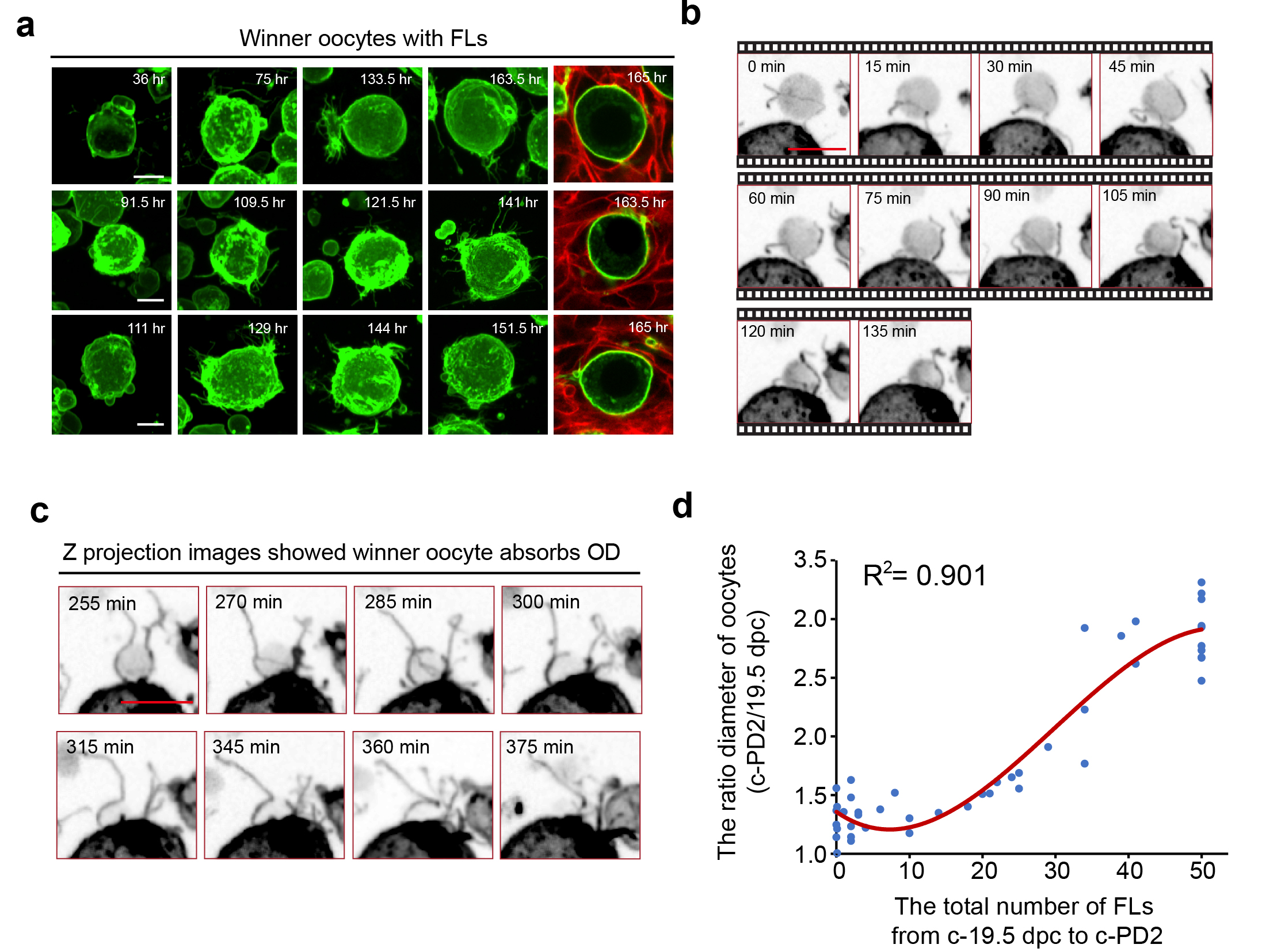

**Extended Data Figure 5. Recording and analyzing the progress of oocyte competition.**

**(a)** Tracing the development of winner-oocytes that form ovarian follicles in culture. Showing the formation of FLs on the winner-oocytes at different time points. **(b-c)** The time-lapse imaging recorded the detail behaviors of oocytes during oocyte competition with 15 interval minus. Imaging the detailed progress of FLs leading OD to attach winner-oocyte (b), and then the winner-oocyte to absorbed the ODs with the help of FLs (c). Oocytes were inverted to black/white (b/w). Scale bar in (a-c): 10 μm. **(d)** The statistical curve showed the positive correlation between the growth ratio of winner oocytes and number of FLs in each oocyte.

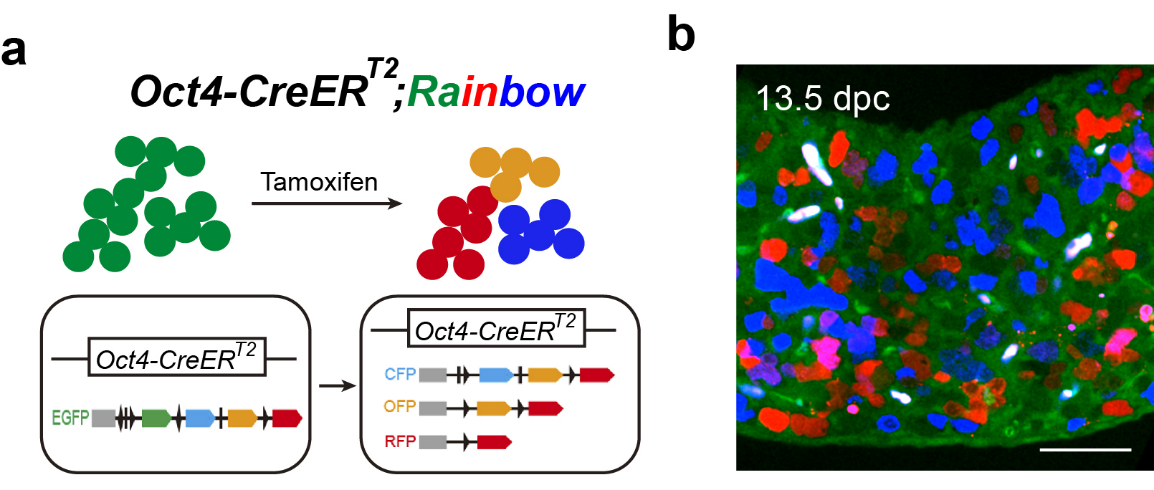

**Extended Data Figure 6. Labeling germ cell cytoplasm in the *Oct4-CreER^T2^;Rainbow* ovaries.**

**(a)** Illustration of tamoxifen (Tam)–induced labeling of germ cells in *Oct4-CreER^T2^;Rainbow* ovaries. In *Oct4*-expressing germ cells, the CreER^T2^ recombinase is not active and the cells express EGFP. Upon tamoxifen injection, the CreER^T2^ recombinase deletes the EGFP region and switches on a random expression of CFP, OFP or RFP. Thus, the cytoplasm of different *Oct4*-expressing germ cells was labeled with blue, orange or red fluorescence. **(b)** The pregnant females with *Oct4-CreER^T2^;Rainbow* fetus were given i.p. injection of 50 mg.kg^-1^ BW tamoxifen at 10.5 dpc, and the labeled fetal ovaries were collected at 13.5 dpc. Scale bar: 50 μm.

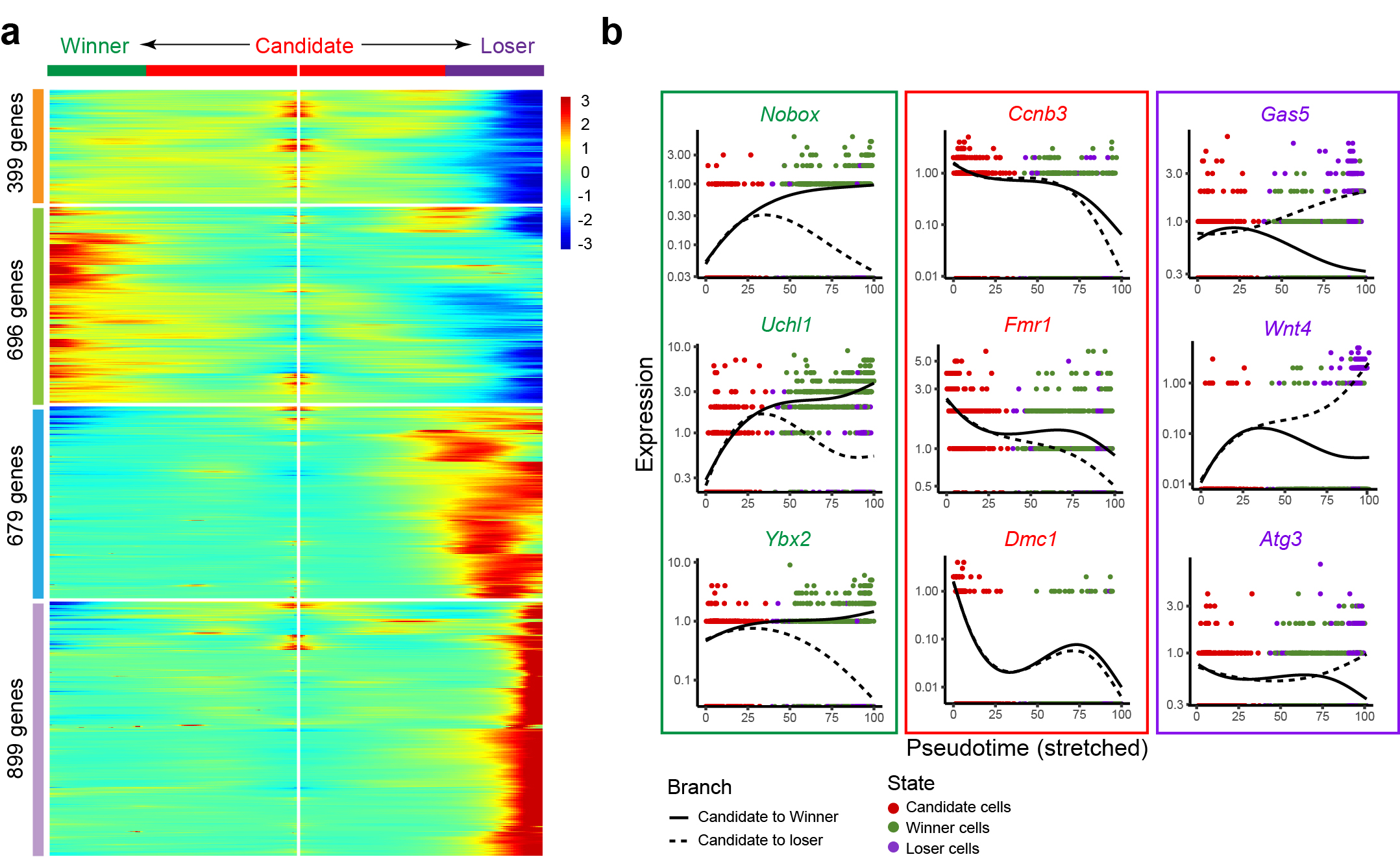

**Extended Data Figure 7. The expression dynamics of the representative genes in the different cell lineages.**

**(a-b)** Heat map (a) and plot (b) of the expression branched dynamics of the representative genes in the different cell lineages in the pseudotime analysis. Expression patterns of the candidate lineage genes *Ccnb3*, *Fmr1*, and *Dmc1*; Winner lineage expressing follicle related genes *Nobox*, *Uchl1*, and *Ybx2*; and Loser lineage expressing genes *Gas5*, *Wnt4*, and *Atg3*.

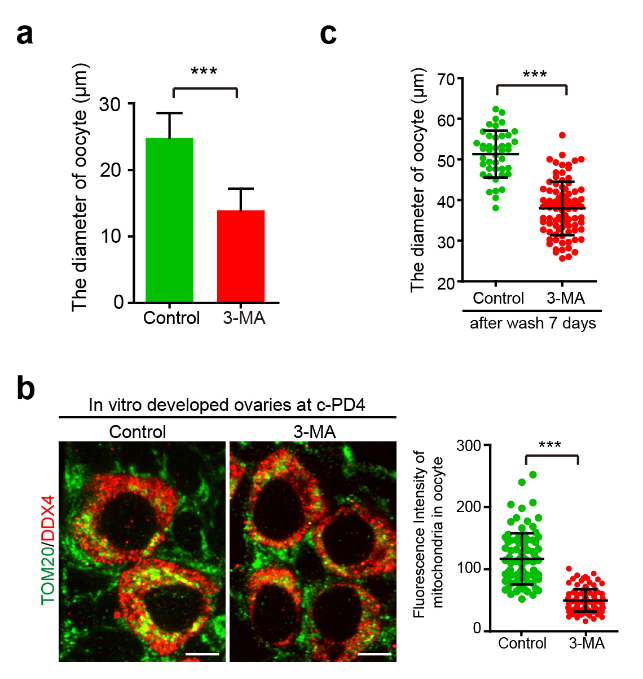

**Extended Data Figure 8. Blocking oocyte competition by 3MA treatment suppressed the growth and the enrichment of mitochondria in the survived oocytes.**

**(a)** Quantification of oocyte average diameter showed significant growth retardation of oocytes in 3-MA treated ovaries compared to those in the control ovaries at c-PD4. Control: n = 48 oocytes, 3-MA: n = 97 oocytes. **(b)** 3-MA treatment decreased the density of mitochondria in survived oocytes. Red: DDX4, Green: TOM20. Scale bar: 10 μm. Control: n = 101 oocytes, 3-MA: n = 106 oocytes. **(c)** Quantification of oocyte average diameter showed oocytes without competition in 3-MA treated ovaries were unable to fully grow. Control: n = 43 oocytes, 3-MA: n = 88 oocytes.
